# Simulations of sequence evolution: how (un)realistic they are and why

**DOI:** 10.1101/2023.07.11.548509

**Authors:** Johanna Trost, Julia Haag, Dimitri Höhler, Laurent Jacob, Alexandros Stamatakis, Bastien Boussau

**Affiliations:** Biometry and Evolutionary Biology laboratory (LBBE) University Claude Bernard Lyon 1, Lyon, France; Computational Molecular Evolution Group, Heidelberg Institute for Theoretical Studies, Heidelberg, Germany; Sorbonne Université, CNRS, IBPS, Laboratory of Computational and Quantitative Biology (LCQB), UMR 7238, Paris 75005, France; Biodiversity Computing Group, Institute of Computer Science, Foundation for Research and Technology - Hellas, Heraklion, Crete, Greece; Institute for Theoretical Informatics, Karlsruhe Institute of Technology, Karlsruhe, Germany

## Abstract

**Motivation:** Simulating Multiple Sequence Alignments (MSAs) using probabilistic models of sequence evolution plays an important role in the evaluation of phylogenetic inference tools, and is crucial to the development of novel learning-based approaches for phylogenetic reconstruction, for instance, neural networks. These models and the resulting simulated data need to be as realistic as possible to be indicative of the performance of the developed tools on empirical data and to ensure that neural networks trained on simulations perform well on empirical data. Over the years, numerous models of evolution have been published with the goal to represent as faithfully as possible the sequence evolution process and thus simulate empirical-like data. In this study, we simulated DNA and protein MSAs under increasingly complex models of evolution with and without insertion/deletion (indel) events using a state-of-the-art sequence simulator. We assessed their realism by quantifying how accurately supervised learning methods are able to predict whether a given MSA is simulated or empirical.

**Results:** Our results show that we can distinguish between empirical and simulated MSAs with high accuracy using two distinct and independently developed classification approaches across all tested models of sequence evolution. Our findings suggest that the current state-of-the-art models fail to accurately replicate several aspects of empirical MSAs, including site-wise rates as well as amino acid and nucleotide composition.

**Data and Code Availability:** All simulated and empirical MSAs, as well as all analysis results, are available at https://cme.h-its.org/exelixis/material/simulation_study.tar.gz. All scripts required to reproduce our results are available at https://github.com/tschuelia/SimulationStudy and https://github.com/JohannaTrost/seqsharp.

## 1 Introduction

Reconstructing the evolutionary history of species or genes by inferring phylogenetic trees is a ubiquitous task in comparative genomics. Typically, phylogenetic inference is based on an MSA that contains aligned sequences of the species under study. A plethora of inference algorithms, tools, and models have been developed to infer phylogenetic trees based on the MSA, for example RAxML-NG [29], IQ-Tree [34], BEAST [9], or RevBayes [24]. When developing novel methods and validating their performance, comparing them to existing state-of-the-art methods on both, empirical, and simulated data is mandatory. Simulated data are particularly useful for conducting inference accuracy and implementation verification assessments, when a known, ground truth phylogeny is needed. Both simulation tools [33, 17, 12] and state-of-the-art inference methods are based on probabilistic models of sequence evolution. Most of the latter exploit models through likelihood functions, by searching for trees that maximize this likelihood [29, 34] or by sampling from posterior distributions via Metropolis-Coupled Markov Chains (MCMC), which also require likelihood computations [9, 24]. Alternatively, researchers have started to explore likelihood-free approaches (for examples outside our field, see Lueckmann et al. [32]). These approaches sample the posterior density instead of evaluating it, and thereby avoid computing the likelihood. The resulting simulated samples are used to build an estimate of the posterior distribution. This so-called simulation-based inference paradigm was pioneered in population genetics under the Approximate Bayesian Inference (ABC) framework [14], and extended over the past decade to neural density estimation techniques [36], where a neural network is trained to output the correct distribution of parameters for a given input observation. In the context of phylogenetic inference, neural density estimation has been restricted to the reconstruction of a single tree as opposed to a full distribution. For example, Suvorov et al. [48] use convolutional neural networks to reconstruct phylogenies from alignments with 4 sequences, and Nesterenko et al. [35] use a transformer-based network architecture to predict evolutionary distances between all pairs of sequences in an alignment.

In all these contexts — evaluation, likelihood-based, or -free inference — it is essential that the probabilistic model of sequence evolution is consistent with empirical data. For evaluation, performance on simulated data is indicative of performance on empirical data, only if the two are sufficiently similar. For inference, a misspecified model can lead to inaccurate and misleading results. For training learning-based methods, it is important that the training data and empirical data are sufficiently similar to circumvent “out-of-distribution” problems [13]. Such problems occur when the training data does not accurately represent the empirical data or when it misses subgroups of the empirical data: the trained method has never “seen” data similar to the empirical data, and can thus behave poorly.

Authors using simulated data in their publications typically set simulation parameters according to attributes (e.g., MSA lengths, or proportions of gaps) of empirical reference MSAs (see e.g., Price et al. [40]). Some also attempt to extract or sample simulation parameters from Maximum Likelihood estimates in large scale empirical databases, such as TreeBASE [38]. The intention is that thereby, simulated data will more closely resemble empirical data [1, 23]. Despite this effort, there still exist performance and/or program behavior differences on simulated versus empirical data. For example, Guindon et al. [20] conclude that comparing methods using simulated data is not sufficient, as “the likelihood landscape tends to be smoother than with real data”, and Hoehler et al. [23] noticed differences between empirical and simulated data when comparing ML phylogenetic inference methods. They conclude that there exist not yet understood differences between simulated and empirical data.

Here, we introduce a metric to quantify how realistic a substitution model is, by simulating data using the respective model and training a classifier to discriminate between simulated and empirical data. We expect realistic models to produce data that are hard to discriminate and induce low classifier accuracy. We leverage recent data simulation tools [33, 17, 12] that are feature-rich and support a wide range of evolutionary models and simulation parameters. We show that we can distinguish simulated from empirical data with up to 99% classification accuracy, depending on the used simulation model. We present two different and independently developed machine learning approaches exploiting distinct MSA characteristics for this classification task: One, using Gradient Boosted Trees (GBT), and another approach based on a Convolutional Neural Network (CNN). We show that prediction accuracy decreases, the more complex the model of evolution used in simulations becomes. Yet, we also observe exceptions to this general trend. For the most complex models in our experimental setup, the prediction accuracy is still very high, with the CNN based classifier achieving prediction accuracies ≥ 0.93 on all tested models. This indicates that simulated alignments are easy to distinguish from empirical alignments, as they do not appear to reproduce some characteristic features of empirical MSAs. We further show that simulating indels remains a challenging task, as including indels results in higher classification accuracies with the CNN classifiers compared to simulations without indels. Further, based on the feature importances of the GBT classifiers, we show that simulated data have more evenly distributed site substitution patterns than empirical data.

## 2 Methods

The goal of our study was to be able to distinguish between empirical and simulated DNA and protein data with high accuracy under increasingly complex models of sequence evolution. Figure 1 depicts our experimental setup for one exemplary set of empirical MSAs (empirical data collection) and one exemplary model of evolution. Using the empirical data collection and the given model of evolution, we simulated a new set of MSAs (simulated data collection) using the AliSim sequence simulator [33]. Based on the empirical and simulated data collections, we completely independently trained two distinct classifiers for each simulated data collection: a Gradient Boosted Tree (GBT) and a Convolutional Neural Network (CNN).

**Figure 1:**
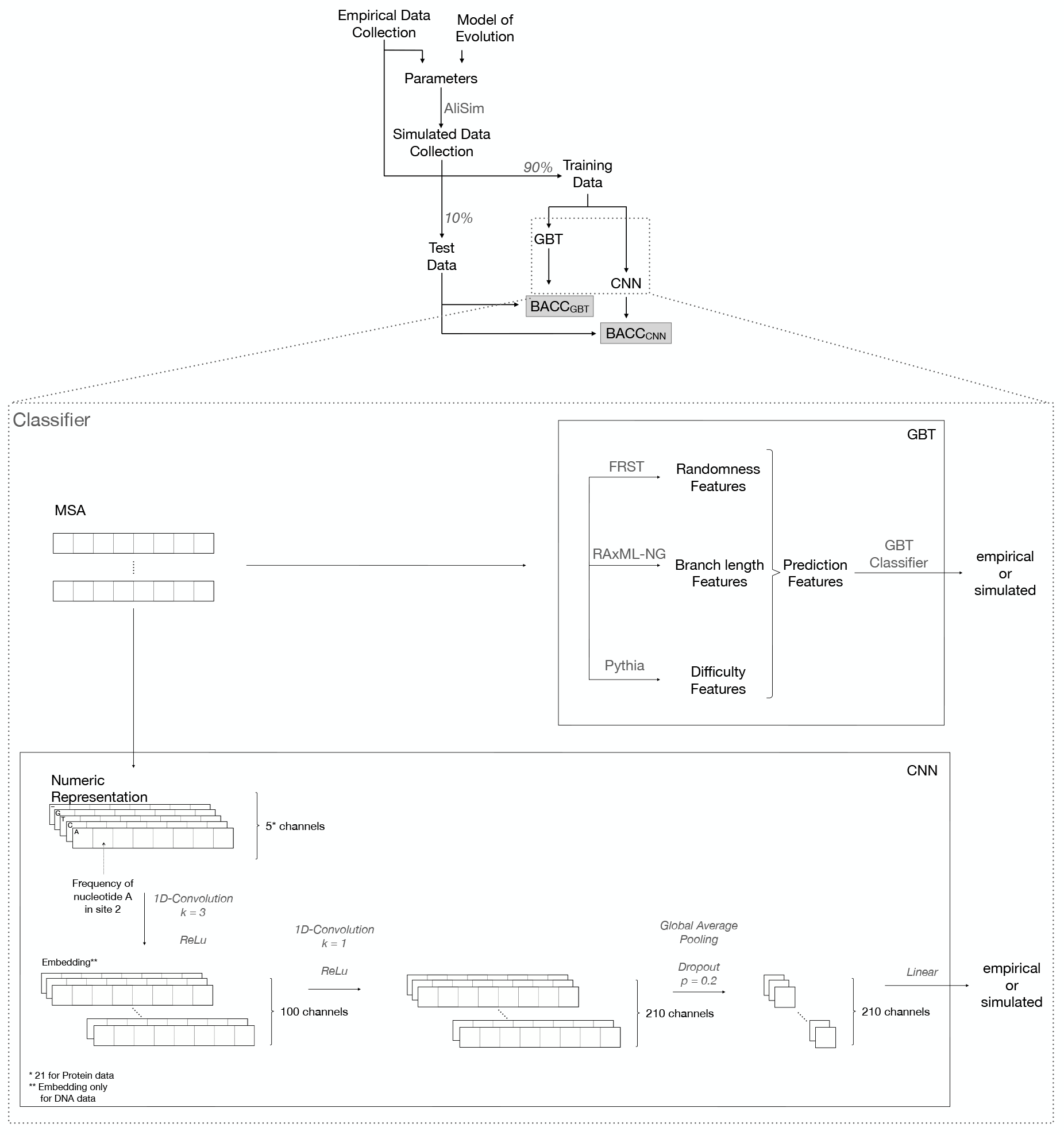
Schematic overview of our experimental setup. Based on a set of empirical MSAs (empirical data collection), we determined parameters for sequence simulation and simulated new MSAs (simulated data collection) under a specific model of evolution using AliSim. Using the empirical and simulated data collections, we trained two distinct classifiers: a Gradient Boosted Tree (GBT) and a Convolutional Neural Network (CNN). The goal of both classifiers is to distinguish empirical from simulated MSAs. For training and evaluating our classifiers, we used a 10-fold cross validation procedure (not depicted for simplicity). In each fold, 90% of the data were used for training and 10% were used for performance evaluation. We evaluated the overall performance of the classifiers via balanced accuracy (BACC).

In the following sections, we describe our experimental setup, the sequence simulation process, and both classification methods in more detail.

### 2.1 Alignment Simulations

For our study, we separately considered DNA and protein data. We simulated 15 MSA sets, seven sets containing DNA MSAs and eight containing protein MSAs, respectively. In the following, we refer to an MSA set as a *data collection*. To simulate the MSAs for each data collection, as well as for data discrimination, we used two empirical data collections as reference, one per data type. The empirical DNA data collection contains MSAs obtained from TreeBASE [38]. The empirical protein data collection consists of MSAs obtained from the HOGENOM database [37]. We removed outliers based on MSA length (i.e., number of sites), number of sequences, as well as MSAs with less than four sequences to ensure a reliable and efficient analysis. Very long sequences would inflate the memory footprint of the CNN, while very short MSAs are uncommon and are more difficult to accurately classify as empirical or simulated. Removing outliers allowed us to deploy a balanced and representative data collection that facilitates robust and unbiased predictions.

Moreover, empirical MSAs may contain sites with ambiguous or exceptional amino acid (AA)/DNA codes, that are ‘B’, ‘Z’, ‘J’, ‘U’, ‘O’ and ‘X’ for protein MSAs, and ‘N’, ‘D’, ‘H’, ‘V’, ‘B’, ‘R’, ‘Y’, ‘K’, ‘M’, ‘S’, ‘W’ and ‘X’ for DNA MSAs. As a further pre-processing step, yet exclusively for the CNN classifier, we removed all MSA sites containing at least one ambiguous letter, as they would bias the prediction. For protein data this concerned 912 out of 6969 MSAs, and we removed 1.34% of all sites within these MSAs. Furthermore, 13.24% of sites in 6117 MSAs with DNA sequences were removed.

For each data type, we generated simulated data collections based on the corresponding empirical data collection, resulting in identical numbers of simulated and empirical alignments. We simulated data using the AliSim sequence simulator [33] under several evolutionary models ranging from easy to complex, in terms of number of free parameters and computational methods used to derive respective AA substitution models. The goal of this setup was to progressively increase simulation realism. First, we simulated five DNA and seven protein data collections without gaps, which allowed us to characterize the realism of substitution models per se. To this end, we removed all sites containing gaps (‘-’) from all empirical MSAs. The resulting empirical data collections contain 7637 DNA MSAs and 6971 protein MSAs, respectively. We henceforth refer to these data collections as *gapless* data collections. Second, we simulated two DNA and one protein data collection with indel events, based on the empirical MSAs containing gaps (9460 DNA MSAs and 6971 protein MSAs).

Note that we chose TreeBASE as the source for empirical DNA alignments, as it is a database of published alignments and thus best represents data that are analyzed in real-world applications of phylogenetics. TreeBASE contains heterogeneous data without a specific focus on the type of underlying genes. See Section 6 in the Supplementary Material for further information on the TreeBASE data.

In the following, we describe the simulation procedures for both data types, as well as our approach to simulate indel events, in more detail. Figures S1 and S2 in the Supplementary Material provide a detailed schematic overview of all simulation procedures.

#### 2.1.1 DNA Simulation

We simulated seven DNA data collections in total (5 gapless and 2 with simulated indel events), with each data collection simulated separately under a different evolutionary model with increasing model complexity. We used the following models of evolution. As the simplest model, we used the Jukes-Cantor (*JC*) model (equal substitution rates and equal base frequencies) [25]. We also used the *HKY* model (four degrees of freedom) [22], and the General Time Reversible (*GTR*) model (eight degrees of freedom) [49]. To account for among site rate heterogeneity, we additionally simulated under GTR in conjunction with the Γ model [53] using four discrete rates (*GTR+G*). The most complex model of evolution we used for simulation was the GTR+G model, with an additional free parameter to accommodate the proportion of invariant sites (*GTR+G+I*) [45].

We selected 9460 empirical MSAs (*Set1*) from TreeBASE [38, 51] as basis for our simulations. We removed all sites containing gaps (‘-’) or fully undetermined characters (‘N’) from the MSAs of *Set1*. Thereby, we obtained 7637 non-empty MSAs (i.e., MSAs that still contained at least one site), which we defined as *Set2*. This lead to an MSA length reduction of around 55% compared to *Set1*. We based our five simulated DNA data collections without indel events on *Set2*, and the two data collections with indels on *Set1*.

AliSim simulates sequences along a given phylogenetic tree. We avoided the problem of simulating realistic phylogenetic trees for this purpose by initially estimating a best-known ML tree using RAxML-NG [29] (default parameters), for every MSA of *Set2* under each of the five evolutionary models (JC, HKY, GTR, GTR+G, GTR+G+I). We then used the inferred phylogeny and respective estimated model parameters to simulate MSAs using AliSim [33] based on every MSA of *Set2*, without specifying an indel model. In the following analyses, we refer to the resulting five gapless data collections as JC, HKY, GTR, GTR+G, and GTR+G+I according to the model of evolution used. In Section 2.1.3 below, we describe the simulation of the two additional DNA data collections with indel events.

#### 2.1.2 Protein Simulation

We simulated seven protein data collections limited to substitution events only, and one additional data collection *with* indels. The most rudimentary evolutionary model we used is the *Poisson* model, with equal exchangeabilities and equal stationary frequencies. We further used two empirical substitution models: the *WAG* [52] and the *LG* [30] model. The LG model is expected to produce more realistic simulations than the WAG model as the former was derived from a larger and more diverse data collection, using more refined inference techniques than the latter. These substitution models use a single set of stationary frequencies (i.e., one AA profile) to simulate all sites in an MSA. We also used mixture models that incorporate heterogeneity among sites by employing multiple profiles. In such models, a profile is drawn from a set of profiles to simulate a single site.

We used the following two mixture models: the C60 model with 60 profiles (*LG+C60*) [46] and the more recent UDM model with 256 profiles (*LG+S256*) [42]. The advantage of the latter model is that each profile is assigned a probability (i.e., weight) of generating a site, while under the C60 model profiles are drawn with equal probabilities. In addition, the UDM model is based on a subset of MSAs from the HOGENOM database, and should therefore generate alignments that are similar to empirical HOGENOM MSAs. To increase model complexity, we performed further simulations accounting for among site heterogeneity using the Γ model [53], as for DNA simulations. We simulated two data collections, one using four discrete Γ rate categories (*LG+S256+G4*) and the second applying a continuous Γ distribution (*LG+S256+GC*).

We set the *α* shape parameters of the Γ distributions based on the values inferred during tree reconstruction when building the HOGENOM database (see Supplementary Material Section 2.1.1). In the following, we will refer to these parameters as *empirical α parameters*. For the simulations, we drew *α* parameters from the probability density function (PDF) estimate of the empirical *α* parameters (see Supplementary Material Section 2.1.2). We compared the ECDF of the empirical *α* parameters with the ECDF of 7000 samples from the PDF estimation to confirm that our distribution of simulated *α* parameters matches the empirical distribution well (see Supplementary Figure S3). We sampled the MSA lengths we used for MSA simulations from the approximated empirical distribution of HOGENOM MSA lengths, using the same approach as for the *α* parameters outlined above. Respective PDF and ECDF functions can also be found in the Supplementary Material in Figure S3. In addition, we compared the AA diversity of empirical protein data and simulations under the LG and LG+S256 models (see Supplementary Figure S6). We simulated all seven data collections along phylogenetic trees that were reconstructed from empirical HOGENOM MSAs where sites containing indels were removed. We performed the inferences using RAxML-NG (see Section 2.1.1).

In analogy to the simulated DNA data collections, we refer to the simulated protein data collections according to the model of evolution used. The gapless protein data collections are Poisson, WAG, LG, LG+C60, LG+S256, LG+S256+G4, and LG+S256+GC. In the following section, we describe the simulation procedure for the data collection with indels.

#### 2.1.3 Simulating Indels

In addition to the gapless data collections, we simulated two DNA, and one protein data collections *with* indels. For both data types, we used the most complex models of evolution as a basis (GTR+G+I for DNA, LG+S256+GC for protein).

To generate the first DNA data collection with indels, we ran tree searches using RAxML-NG under the GTR+G+I model for each MSA of DNA *Set1*. We then simulated MSAs with indels using two distinct procedures to generate two distinct data collections. For the first data collection, we simulated the MSAs in the same way as for the gapless collections. Then, we superimposed the gap pattern of the MSAs used as the basis of the simulation onto the simulated MSAs. We refer to this procedure as the *mimick* procedure and denote the resulting data collection as GTR+G+I+mimick.

For the second data collection, as well as the protein data collection with indels, we simulated the MSAs using not only the inferred trees and estimated evolutionary model parameters, but also specifying indel parameters. In the following, we describe the procedure to infer and validate these parameters. We performed this procedure for both DNA and protein data collections separately. We refer to this procedure as the *sparta* procedure. We first used the SpartaABC tool [31] to obtain indel-specific parameters from a subset of empirical MSAs. Here, we employed the rich indel model (RIM), which differentiates between insertion and deletion events using five free parameters. The inferred parameters are: Insertion and deletion rate (I R, D R), root length (RL), and the parameter *a* that controls the Zipfian distribution of insertion and deletion lengths (A I, A D). We will henceforth refer to this set of parameters as *empirical indel parameters*.

To simulate MSAs, we drew indel parameters from the joint parameter distribution of empirical indel parameters. To approximate the probability density function (PDF), we applied Gaussian kernels to the five principal components of the indel parameters. This choice was based on our observation that a more accurate match is achieved between the empirical parameters’ empirical cumulative distribution function (ECDF) and the resulting parameters’ ECDF when using the principal components. For the Gaussian kernels, we determined the bandwidth using Scott’s rule of thumb [43]. Moreover, we employed the kernel-density estimation implementation by Virtanen et al. [50], although it tends to overestimate the distribution’s actual edges. To mitigate this issue, we re-sampled values if they fell outside the bounds of the parameter prior bounds chosen by Loewenthal et al. [31]. To validate our approach, we compared the ECDF of the empirical parameter values with the ECDF of parameters sampled from the empirical PDF for each indel parameter type. Plots of the ECDFs and density functions are provided in the Supplementary Material Figure S4 and Figure S7. Moreover, we compared the density functions of empirical and simulated MSA lengths as a sanity check (see Figure S5 and Figure S8 in the Supplementary Material). We denote the resulting DNA data collection as GTR+G+I+sparta, and the resulting protein data collection as LG+S256+GC+sparta.

### 2.2 Classification Methods

To distinguish simulated and empirical MSAs, we developed two distinct approaches. One approach is a standard machine learning algorithm based on hand-crafted features and Gradient Boosted Trees (GBT). Using GBTs allows us to attain insights on feature importance, explain the classification results, and determine short-comings of MSA simulations. Our second approach uses Convolutional Neural Networks (CNN). In contrast to GBT, CNNs only require minimal data processing as they are able to automatically learn relevant features through training. However, to interpret these features, additional analysis is necessary. In the following, we introduce both machine-learning approaches to classification, and describe our training setups.

### 2.3 Training classifiers

In this section, we briefly describe how we trained our classifiers and introduce useful terms for readers that are unfamiliar with machine learning. Classifiers are functions that, in our case, take as input a MSA or MSA features and output the label “simulated” or “empirical”. These functions depend on numerous parameters, whose values must be set during a so-called learning phase. This learning phase uses MSAs annotated with their ground truth labels. The function is applied to each alignment, and its predicted label is compared to the true label, thanks to a cost function that is also often called loss function. Parameter values are then refined based on the computed cost. By iteratively going through each alignment, the parameter values are tuned, and the accuracy of the classifier typically improves. When all the training alignments have been examined by the function once, we say that an epoch has passed. The training is stopped after a number of epochs, typically because the number of iterations has been limited a priori (e.g., for the GBT classifier), or because classifier accuracy does not improve further (e.g., for the CNN classifier). To assess the performance of a classifier after the training, it is important to use data that has not been part of the training data set. For this reason, we split our alignment collection into two categories: most of the alignments were used for training (training data, 90% of all MSAs), and a subset was used for evaluating the performance of the classifier (test data, 10% of all MSAs). We repeated the training and evaluation 10 times, on different random splits of the data (*i*.*e*., 10-fold cross validation), and averaged over the respective 10 performance/accuracy metrics. We used the balanced accuracy metric (BACC) [11] to assess performance, as this metric allows for varying proportions of simulated/empirical MSAs in the data collection and better reflects classification accuracy for imbalanced datasets. The balanced accuracy is the average of the sensitivity (here, 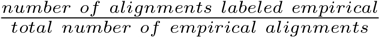) and specificity (here, 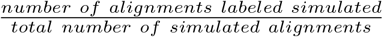). The best BACC value is 1 and the worst value is 0.

#### 2.3.1 Gradient Boosted Trees

Gradient Boosted Trees (GBT) is an ensemble machine learning technique that combines multiple decision trees to obtain an accurate prediction model [18]. Training a GBT classifier consists of *M* sequential stages, with each stage contributing an additional decision tree that improves the estimator of the previous stage. For our experiments, we used the GBT classifier as implemented in the LightGBM framework [26].

##### Prediction Features

To classify MSAs into simulated or empirical ones, we computed 23 features for each MSA. Four of these features are attributes of the MSA: the *sites-over-taxa ratio*, the *patterns-over-taxa ratio*, the *patterns-over-sites ratio* and the proportion of invariant sites (*% invariant*). For data collections simulating indel events, we also used the proportion of gaps as feature (*% gaps*). Further, we quantified the signal in the MSA using the difficulty of the respective phylogenetic analysis as predicted by Pythia [21] (*difficulty*), as well as the Shannon entropy [44] of the MSA (*Entropy*), a multinomial test statistic of the MSA (*Bollback multinomial*; [8]), and an entropy-like metric based on the number and frequency of patterns in the MSA (*Pattern entropy*). For further details on the computation of these metrics, we refer the interested reader to Supplementary Material Section 4.1. In order to assess downstream effects on tree inferences using simulated and empirical data, we inferred 100 trees based on the fast-to-compute maximum parsimony criterion [15, 16] and a single Maximum Likelihood (ML) tree using RAxML-NG [29]. We added two features based on the inferred 100 maximum parsimony trees: the average pairwise topological distance using the Robinson-Foulds distance metric (*parsimony RF-Distance*) [41], as well as the proportion of unique topologies (*% parsimony unique*). We further refer to these features as difficulty features. Based on the ML tree inferred by RAxML-NG, we computed a set of branch length features, namely the minimum, maximum, average, standard deviation, median, and sum of all branch lengths in the ML tree (*brlen*_*min*_, *brlen*_*max*_, *brlen*_*avg*_, *brlen*_*std*_, *brlen*_*med*_, *brlen*_*sum*_).

We used the next six features to highlight one of the recurrent problems of simulated sequence generators: a common simplification used in generators is the assumption that substitutions occur at uniformly distributed random locations in the sequence, which appears to not be the case in real-world genetic data [10]. Thus, we expected empirical MSAs to be less uniform than simulated MSAs, and we henceforth attempted to confirm this hypothesis.

To quantify substitution frequency distributions along an MSA, we first inferred a parsimony tree using RAxML-NG. Then, based on the parsimony criterion, we calculated the number of substitutions for every site, resulting in a vector *m*. Given the vectors *m* for empirical and simulated MSAs, we can anecdotally observe that the locations of substitution occurrences appear to be less uniformly distributed in empirical than in simulated MSAs (see Figure 2, more examples available in Supplement Material Section 4.3). To the best of our knowledge, there is no panacea in quantifying the absence of structure in data, and it is part of ongoing research in the field of cryptography. We resorted to the Fourmilab Random Sequence Tester (FRST) (https://www.fourmilab.ch/random/), that is used to evaluate pseudo-random number generators, to quantify randomness in *m*. FRST computes six measures of randomness: Entropy (*Entropy*_*rand*_), maximum compression size reduction in percent (*comp*), Chi-Square test (*Chi*^2^), arithmetic mean (*mean*_*rand*_), Monte Carlo Value for Pi (*mcpi*) (see Section 4.2 in Supplementary Material), and Serial Correlation Coefficient (*SCC*) [28]. We executed FRST with a binary representation of *m* on all data collections, then we normalized the computed measures of randomness, and used these values in our predictions. We henceforth refer to this set of six features as *randomness features*.

**Figure 2:**
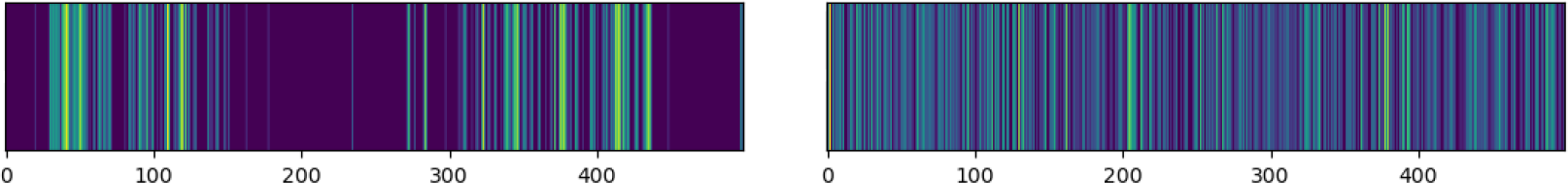
Visualized substitution rates for an anecdotal (specifically selected to highlight the issue) gapless empirical DNA MSA (left), and gapless simulated MSA (right) generated based on the inferred tree and estimated evolutionary model parameters of the left MSA under the GTR model. The x-axis denotes the alignment site index. A brighter color denotes a higher number of substitutions.

##### Training and Optimization

For each of the simulated data collections presented above, we trained a distinct binary GBT classifier. We trained each GBT classifier using a stratified 10-fold cross-validation procedure. Here, stratified means that the proportion of empirical and simulated MSAs in both training and test subsets was the same. The training data consisted of one simulated data collection and the empirical data collection for the respective data type. We used the hyperparameter optimization framework Optuna [4] to determine the optimal set of hyperparameters for each classifier. For each GBT classifier, we performed 100 Optuna iterations using a Tree-structured Parzen Estimator algorithm [7] to sample the hyperparameter space. To prevent the classifiers from overfitting the data, based on preliminary experiments, we limited the depth of the individual decision trees to a maximum of 10, the maximum number of leaves to 20, and the minimal number of samples per leaf to 30. Additionally, we applied L1 and L2 regularization to prevent overfitting and better generalize to unseen data [19]. We determined the optimal weights of L1 and L2 regularization independently using Optuna. L1 regularization sums over all absolute weights in the decision tree and thus penalizes trees with a high number of branching events. As a result, L1 regularization sets the least important feature weights to 0 and thus selects the most important features for classification, leading to shallower trees. In contrast, L2 penalizes large weights by summing over the *square* of all weights, thereby leading to close to zero weights and thus preventing the classifier to heavily rely on but a few features. A more detailed description of the feature generation, and training process, the hyperparameter optimization, as well as the hardware setup are available in the Supplementary Material Section 4.1.

#### 2.3.2 Convolutional Neural Networks

Convolutional Neural Networks (CNNs) are a popular prediction method originally developed for computer vision and image processing. Recently, they have been applied to predict properties of biological sequences [6, 5, 54]. A CNN jointly learns a representation of the data (through convolution layer(s)) and the classification of the data based on these representations (in our case using a fully connected layer). More precisely, a convolution layer slides short probabilistic sequence motifs along the sequence, and outputs an activation profile (i.e., feature map) for each of these motifs. A motif is called kernel and the length of the motif, kernel size. Here, we used a CNN to classify empirical and simulated MSAs. In the following we will detail the input to the network, its architecture, training and optimization, and the evaluation of its performance.

##### MSA Representation

In order to obtain a numeric representation of an MSA, where the network is invariant to the order of sequences, we used a two-step approach. First we decided to represent the MSA using its site-wise AA or nucleotide composition, i.e., the AA or nucleotide proportions per site, which sum to one. Second, each AA/nucleotide, as well as gaps are passed to the convolution network as input features (i.e., channels), resulting in 5 (4 DNA sites + gap) or 21 (20 AAs + gap) channels. This is analogous to using color channels in an image. It maintains the identity of a nucleotide/AA and is common practice when applying CNNs to biological sequences [6]. The input size was the maximum MSA length in the simulated and empirical data collection. All MSAs with fewer sites were zero-padded at their edges in order to match the fixed input size.

Empirical protein sequences typically start with Methionine (M), which simulations do not account for. We removed the first and second sites from the empirical protein data to avoid biasing the prediction. To evaluate the impact of removing the second site, we tested the trained network on empirical validation data, including the second site. The absolute accuracy difference between data with and without the second site was below 0.0005 (see Table S2 in the Supplementary Material).

##### CNN Architecture

We developed two architectures, one for each data type (DNA and protein). We explored alternative architectures and chose the architecture with the best balance between complexity and performance. For protein MSAs, we used a single one-dimensional convolution layer with 210 filters of size 1 *×* 21 (i.e., kernel size *×* input channels). Of note, these filters do not take into account the phylogenetic structure of the data, and simply capture AA profiles at single sites, as opposed to larger motifs spanning several contiguous sites typically used in CNNs. For DNA sequences, we used a two-layer CNN, whose first layer has 100 filters of size 3 *×* 5 and is meant to capture codon structure. The second layer has 210 filters of size 1 *×*100. A standard Rectified Linear Unit (ReLU) activation function is employed in both architectures [3]. An activation function is a nonlinear transformation of a node’s output. It is applied before passing the output to the next layer. The ReLU outputs its input if it is positive, and zero, otherwise. For both DNA and protein architectures, the layers following convolution comprise a dropout layer, which deactivates a node with a certain probability (here we chose 0.2) to avoid overfitting and global average pooling along the sequences. A final fully-connected layer combines all features (i.e., channels) for the binary prediction. For this, we used a Sigmoid activation function. In total, the protein network counts 4831 learnable parameters, while the DNA network has 23,021 due to the additional convolution layer.

##### Training and Optimization

To update the network weights, we employed the Adam optimizer [27] along with a binary cross-entropy loss function. The optimized parameters include the learning rate and the number of filters. For the former, we chose the learning rate range independently for each data collection and fold using the learning rate range test (LRRT) [47]. The LRRT involves gradually increasing the learning rate during a few training epochs, monitoring the change of the loss, and plotting the results. It helps to select a learning rate where the model effectively learns and quickly converges without extensive manual tuning. Given the LRRT results, we then evaluated different learning rates by means of the validation loss after 100 epochs and considered the learning curve, that is, the validation and training loss over epochs. Furthermore, we varied the number of filters and chose the number that yielded the maximal validation BACC. For more details on the optimized parameters and hardware used for training, see Section 3 in the Supplementary Material. In addition to the validation BACC, we considered the Class Mean Absolute Error (MAE), which is the mean absolute difference between the accuracy on simulated and empirical data collections across folds, as well as the standard error (SE), which denotes the standard error of the obtained validation BACC across folds. If these measures were strikingly large, we interpreted this as an indicator that the network needs to be improved to generalize better. As with the other classifier, we used 10-fold cross-validation. We applied an early stopping rule [39] to automatically terminate the training of every fold individually. However, for certain data collections, we observed that the chosen stopping rule seemed overly strict. The visualized learning curves indicated that the network had converged, even though the stopping criterion was not met. Consequently, we decided to manually terminate the training in these cases. Learning curves, Class MAE, and the SE can be found in the Supplementary Material (Table S1, Figures S9 and S10).

To compare the performance of CNNs trained on various simulated data collections, we determined the maximum validation BACC over training epochs for each CNN and for each fold. What is referred to as BACC is the average BACC across folds at the selected epochs. Because we are using the same validation data to choose the stopping epoch and assess the resulting accuracy, there is a risk that this accuracy is overoptimistic. To quantify this risk, we computed summary statistics of BACCs of epochs surrounding the selected epoch (see Table S1 in the Supplementary Material).

#### 2.3.3 Performance Evaluation

Using the BACC metric per data collection, we compared the performance of pairs of classifiers of simulated data collections. In order to evaluate whether the difference of the BACCs of two data collections and therefore two different evolutionary models is significant, we conducted multiple unpaired two-samples t-tests, where one sample consists of the validation BACC for each fold. This allowed us to compare models in their ability to produce simulations that are more or less or equally realistic. For protein data, we compared the BACCs of the following groups: Poisson vs. WAG, WAG vs. LG, LG vs. LG+C60, and all pair-wise combinations of site heterogeneous models. The null hypothesis is that these models yield equal average BACCs across folds. We rejected the null hypothesis if the resulting P-value was below the significance level of 0.05. For DNA data, we compared the BACCs of JC vs. HKY, HKY vs. GTR, GTR vs. GTR+G and GTR+G vs. GTR+G+I. To account for multiple testing, we applied Bonferroni correction, i.e., we multiplied each P-value by the number of tests for each data type separately [2]. An overview of all tests is provided in the Supplementary Material (Tables S7 to S9).

## 3 Results

Table 1 shows the BACC for our GBT and CNN classifiers across all data collections. Both classifiers were able to accurately distinguish simulated from empirical data. The GBT classifiers achieved high BACCs for all simulated protein data collections (*≥*0.98), as well as for all gapless DNA data collections (*≥*0.89). We observed the worst BACC of 0.77 for the DNA data collection simulated under GTR+G+I with gaps simulated according to the *mimick* procedure. The CNN classifiers achieved BACCs ranging from 0.93 to 0.9996. Interestingly, the GBT classifiers showed similar BACCs or even outperformed the CNN on the protein data collections, but achieved lower BACCs on DNA collections.

**Table 1:** Average of the BACC on empirical and simulated data collections across 10 folds for the GBT and CNN classifiers. Parameter configurations of simulations listed in the first column are sorted with increasing complexity from top to bottom for both DNA and protein data. For both, the last row(s) shows results on data collections with indels.

| Data collection | BACC |  |
| --- | --- | --- |
|  | GBT | CNN |
| DNA data collections |  |  |
| JC | 0.96 | 0.99 |
| HKY | 0.96 | 0.99 |
| GTR | 0.94 | 0.93 |
| GTR+G | 0.89 | 0.94 |
| GTR+G+I | 0.89 | 0.94 |
| GTR+G+I+mimick | 0.77 | 0.97 |
| GTR+G+I+sparta | 0.94 | 0.97 |
| Protein data collections |  |  |
| Poisson | 0.99 | 0.9996 |
| WAG | 0.99 | 0.97 |
| LG | 0.99 | 0.95 |
| LG+C60 | 0.98 | 0.99 |
| LG+S256 | 0.99 | 0.995 |
| LG+S256+G4 | 0.99 | 0.99 |
| LG+S256+GC | 0.98 | 0.99 |
| LG+S256+GC+sparta | 0.99 | 0.996 |

On DNA data collections, substitution models with fewer degrees of freedom than the GTR model, namely JC and HKY, were classified more accurately (BACC=0.99 for CNN and BACC=0.96 for GBT). However, increases in model complexity did not always translate into improvements in the realism of the data. For instance, the performance of the CNN was marginally better on simulations under the HKY model than on simulations under the simpler JC model (P=0.03, see Table S1 in the Supplementary Material). The GBT predictions, which were equally accurate for JC and HKY simulations (BACC=0.96), did not reflect any improvement in the simulations due to more degrees of freedom in the HKY model either. Moreover, the CNN yielded the lowest BACC (0.93) on simulations conducted under the GTR model. In contrast, simulations that included rate heterogeneity (GTR+G) were slightly easier to classify (BACC=0.94, P=0.04). Contrary to our expectations, including a proportion of invariant sites (GTR+G+I) did not result in a lower BACC compared to GTR+G simulations (BACC=0.94, P=1.0 for CNN, BACC=0.89, P=1.0 for GBT).

We did not observe the expected trend of an increased realism with an increase in model complexity for the protein data collections. For instance, the CNN had the lowest BACC on simulations under the LG substitution model (BACC=0.95) and not on the more complex mixture models. For the GBT, distinguishing the LG+S256+G4 data collection appeared to be easier than the data collection based on the simpler LG+C60 model (P=0.77). Unexpectedly, all simulations using a mixture of stationary frequency profiles (i.e., LG+C60, LG+S256, LG+S256+G4 and LG+S256+GC) were nearly perfectly discriminated from the empirical data collection with both GBT and CNN (BACC *≥* 0.98). With the CNN, we did not find a significant performance difference between these evolutionary models (P 0.*≥* 38, see Table S9 in the Supplementary Material).

To rule out the possibility that these rather unexpected findings are a consequence of specific behaviors inherent to the AliSim simulator, we conducted an experiment to evaluate the performance of the CNN classifier pre-trained with LG+S256 simulations on data generated using a simulator developed in house that employs the same model. Our results showed that the CNN classifier performed comparably well on the alternative simulations (BACC=0.99). In addition, we tested the same CNN on simulations using 4096 profiles. These simulations were only slightly harder to classify (BACC=0.98) than the ones based on only 256 profiles (BACC=0.995).

The CNN trained on empirical data collections with indels and simulations under the most complex evolution model with indels (i.e., LG+S256+GC+sparta, GTR+G+I+mimick, GTR+G+I+sparta) also yielded highly accurate predictions (BACC=0.996 for protein and BACC*>*0.97 for DNA data). The results were similar to or better than the results obtained without indels. There was no significant difference between CNN performance on the two DNA indel models employed (P=1.0). Simulating indels increased the GBT classification accuracy for protein data (BACC=0.99) and the *sparta* based DNA data collection (GTR+G+I+sparta; BACC=0.94) compared to the same model of evolution without indel simulations (LG+S256+GC BACC=0.98; GTR+G+I BACC=0.89). We did, however, observe a significant decrease in accuracy comparing the two DNA indel models (P=0.0). GBT classified the GTR+G+I+sparta data collection with high accuracy (BACC=0.94), but showed an unexpectedly low BACC of 0.77 for GTR+G+I+mimick.

In order to gain insights into why the general classification task achieved high prediction accuracy and appears to be rather easy in general, we assessed the influence of the described features on the prediction of the GBT classifiers. To this end, we computed the gain-based feature importance. The gain-based feature importance directly measures the contribution of a feature to the reduction of the loss function. Table S4 in the Supplementary Material shows the three most important features for all classifiers.

We observed that, except for one data collection, the SCC randomness metric was the most important feature. For classifying the LG+S256+GC+sparta data collection, it was the second most important feature. Figure 3 shows the distribution of SCC values for one example DNA data collection (GTR+G+I), as well as for one example protein data collection (LG+S256+GC) compared to the distribution for the respective empirical data collection. The lower the SCC value, the more random is the distribution of rates of evolution across sites in the MSA. The SCC values for simulated MSAs are substantially lower than for empirical MSAs. This shows that the rates of evolution across sites are more uniformly distributed in simulated MSAs compared to empirical MSAs, simulated data is thus more “random” than empirical data. We observed similar patterns for all other data collections as well.

**Figure 3:**
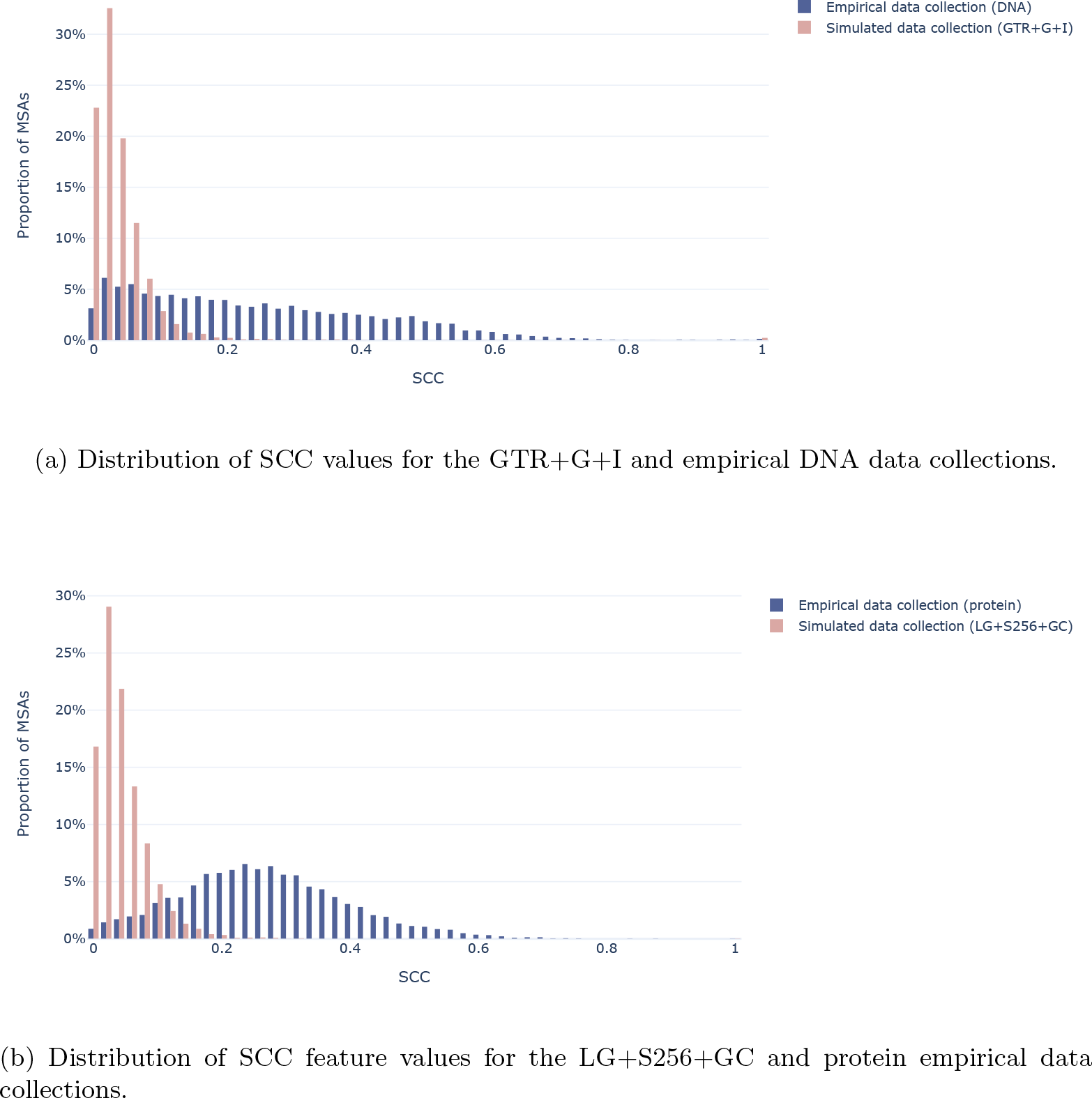
Feature distribution of SCC feature values for one exemplary DNA and protein data collection. The dark blue bars represent the respective empirical data collection and the light pink bars represent the respective simulated data collection.

We also frequently observed the *Entropy*, the *Pattern entropy*, as well as the *Bollback multinomial* metrics being among the three most important features. While the randomness features measure the randomness across *sites* of the MSA, these three features quantify the randomness across *taxa* per site, indicating that simulated data is not only more “random” across sites, but also within sites.

To gain further insights into the importance of the randomness features for classification, we additionally retrained all GBT classifiers without this set of randomness features. Table S6 in the Supplementary Material shows the resulting BACCs alongside the three most important features. As expected, the BACCs decrease for all data collections. Interestingly, the BACCs for the GTR+G and GTR+G+I DNA data collections decreased substantially from 0.89 to 0.65 and 0.61 respectively, yielding a prediction only marginally better than random guessing. Using this reduced set of features for the prediction, we observed interesting differences in feature distributions. We observed that, compared to simulated data, empirical data tends to have a higher proportion of invariant sites (Figure 4a). The branch lengths in trees inferred for simulated MSAs tend to be shorter (Figure 4b; for better visualization, we only show data between the 10% and 90% percentile), and the *parsimony RF-Distance* tends to be higher for empirical data (Figure 4c). While Figure 4 depicts the distribution of feature values for one exemplary data collection (JC) only, these observations hold true for all simulated data collections. The more complex the model of evolution, the less pronounced these differences are, especially for the simulated DNA data under GTR+G and GTR+G+I (see Supplementary Material Figures S15 and S16). It is noteworthy however that even GTR+G+I, which contains a parameter dedicated to modelling the proportion of invariable sites, produces alignments with fewer invariant sites than in empirical data.

**Figure 4:**
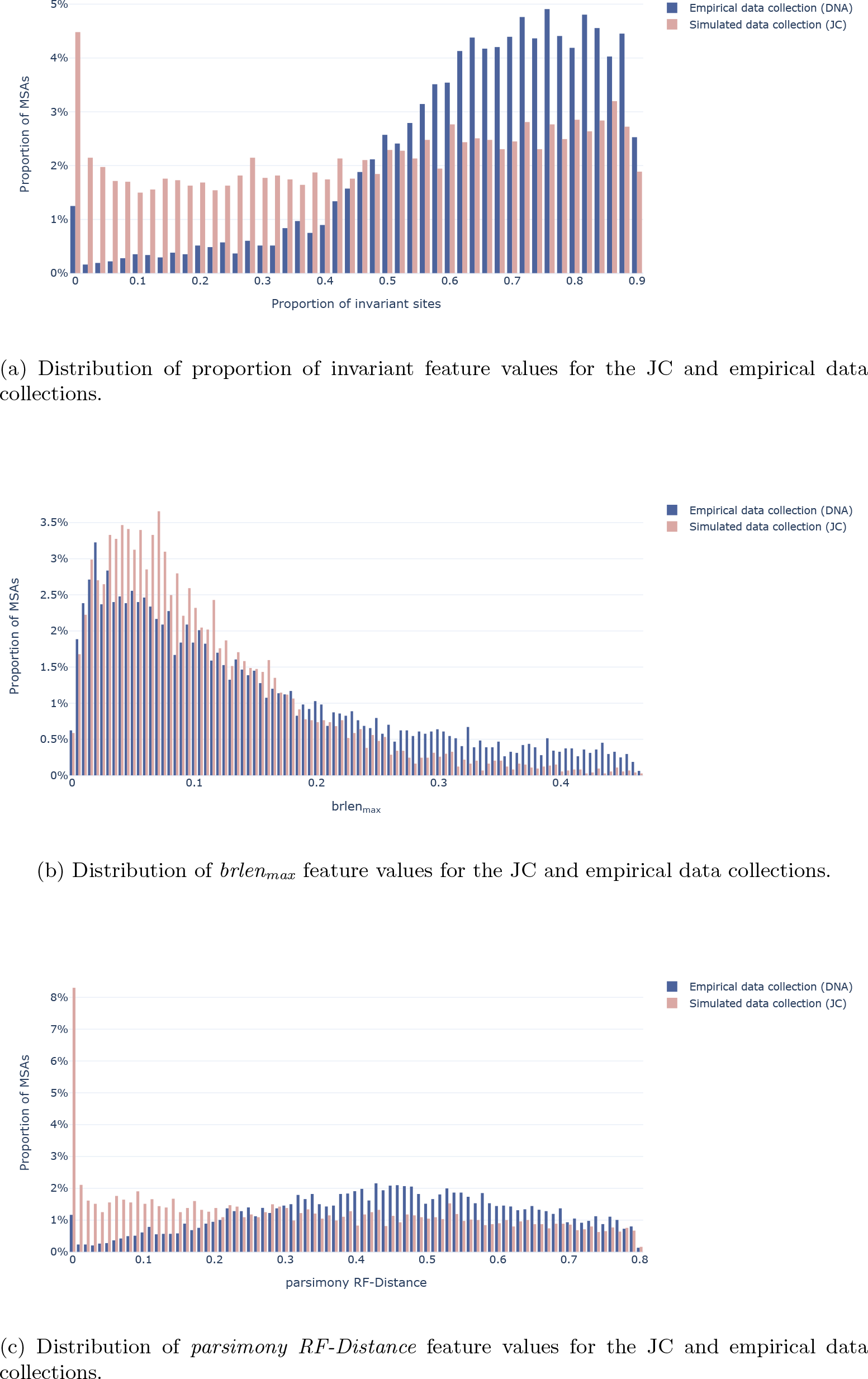
Feature distribution for important features for classifying the JC data collection. The dark blue bars represent the empirical data collection and the light pink bars represent the simulated JC data collection.

We further explored the substantial decrease in accuracy for the GTR+G+I+mimick data with a BACC of 0.77. To this end, we split MSAs site-wise into 100 parts (buckets), averaged the number of substitutions per bucket (normalized by the maximum number of substitutions per MSA), and averaged the buckets over every MSA (see Figures S12 to S14 in the Supplementary Material). Interestingly, we could observe that the substitutions for empirical and the GTR+G+I+mimick data collections are concentrated at the beginning and the end of the MSAs, while the number of substitutions in GTR+G+I+sparta seem to be uniformly distributed. This also seemed to be the case for other substitution models (results not shown). This result is in agreement with Bricout et al. [10] who also found this pattern in a large scale analysis of empirical alignments.

As described above, we simulated the DNA data collections and the protein data collections without indels based on trees inferred using RAxML-NG. Trees for protein data with indels used for our indel simulations were inferred using IQ-Tree. For 10 out of 15 data collections, one of the branch length features was among the three most important features. To ensure that we did not leverage a tool-induced bias for our prediction, we retrained all classifiers using only the MSA-based features by discarding all branch length features. We observed no substantial impact on the overall prediction accuracies. With GTR+G+I+mimick we observed the highest BACC difference. Using all features, the GBT achieved a prediction accuracy of 0.77. Discarding the branch length features resulted in a BACC of 0.74. Table S5 in the Supplementary Material shows the resulting BACCs for all classifiers, alongside the three most important prediction features when only using MSA based features.

In addition to the feature analysis of the GBTs, we further investigated the remarkably accurate performance of the CNN on simulations using mixtures of stationary frequency profiles (i.e., the S256 or C60 model). Given that we could achieve better performance when using average global pooling, that is, averaging across the sequence, instead of maximum local pooling following the convolution layer (see paragraph CNN Architecture) we hypothesized that there must be predictive global features that aid in distinguishing simulated from empirical MSAs. In particular, we hypothesized that alignment-wise frequencies of AAs or nucleotides may differ between simulated and empirical data. To test this hypothesis, we trained logistic regression models to undertake the same classification task, but using site compositions averaged along the alignment, i.e., MSA compositions. Figure 5 shows that the logistic regression model indeed performed well, particularly for simulated data under mixture models (BACC*>*0.94). Moreover, across collections, there is a strong correlation between BACCs of the CNNs and the logistic regression models (*r*^2^ = 0.85). We also attempted to train the logistic regression model on DNA data simulated under the GTR+G+I model, but found that there was no significant improvement during the first 100 epochs (BACC=0.51). Therefore, the MSA composition is not informative for the classification of DNA data, but highly informative for protein data.

**Figure 5:**
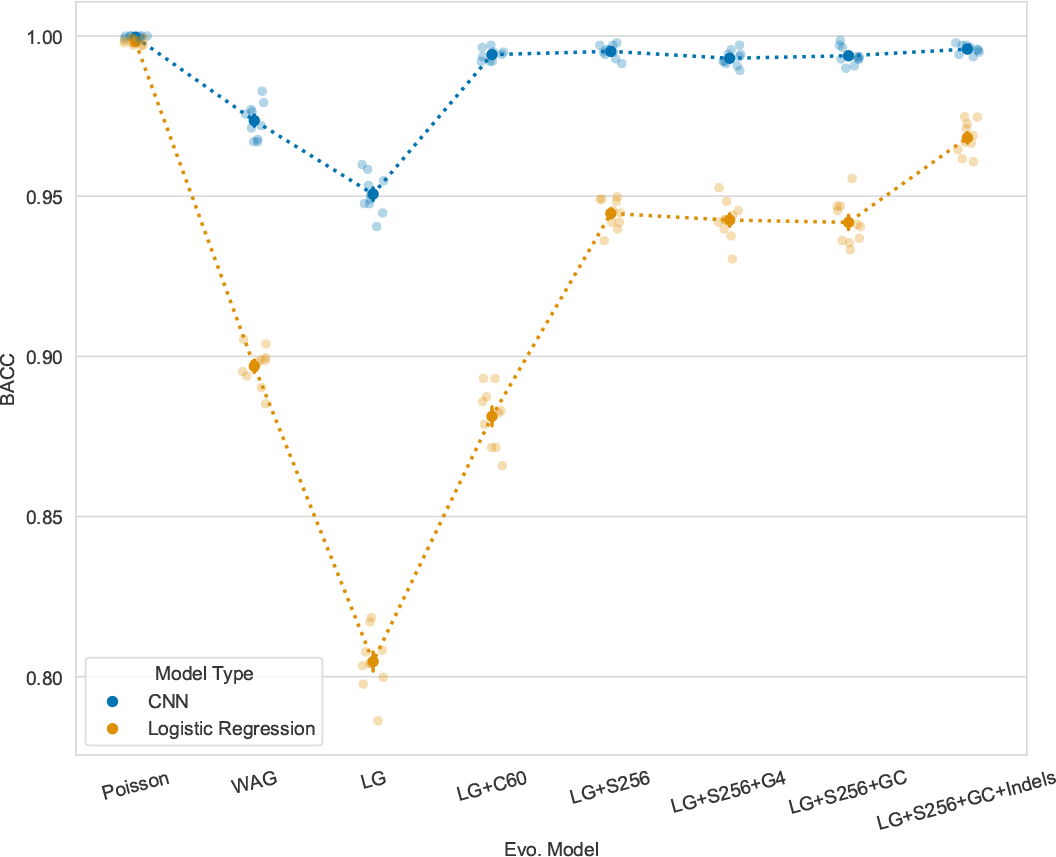
Performance of logistic regression on MSA compositions and CNN on site-wise compositions. For each evolutionary model the BACC of each fold is represented as well as the mean and standard error.

For both GBT and CNN classifiers, we observed a general trend for lower classification accuracy on more difficult MSAs according to the Pythia difficulty score. The higher the Pythia difficulty for an MSA, the lower the signal in the data and the more difficult it is to obtain a well-supported phylogeny as the likelihood surface exhibits multiple indistinguishable (by means of standard phylogenetic significance tests) likelihood peaks [21]. In addition to assessing the BACC as a function of the difficulty of *simulated* MSAs, we also assessed the BACC as a function of the difficulty of the underlying *empirical* MSAs. For MSAs with a higher Pythia difficulty, it is more difficult to obtain a well-supported phylogeny, as the likelihood surface exhibits multiple peaks. However, simulating an MSA requires a reference phylogeny and relying on a “bad” tree might have a negative impact on the realism of the simulated data. If this holds true, we expected the classification of simulated MSAs based on easy empirical MSAs (i.e. simulations based on “good” trees) to be more difficult, leading to a lower BACC than the classification of simulated MSAs based on difficult empirical MSAs. Interestingly, we observed the opposite effect: the more difficult the underlying empirical MSAs, the lower the BACC. Figure 6 depict this observation for the simulated data collections with the lowest BACC for GBT (GTR+G+I+mimick) and CNN (LG) respectively. Both Figures show the BACC as a function of the Pythia difficulty over the *simulated* MSAs (left subfigures), as well as the BACC as a function of the Pythia difficulty over the underlying *empirical* MSAs (right subfigures). We suspect that both observations are related to the amount of signal in the data: MSAs with a low signal not only lead to inconclusive phylogenetic analyses (as indicated by the high Pythia difficulty), but apparently also lack a strong signal that indicates their realism.

**Figure 6:**
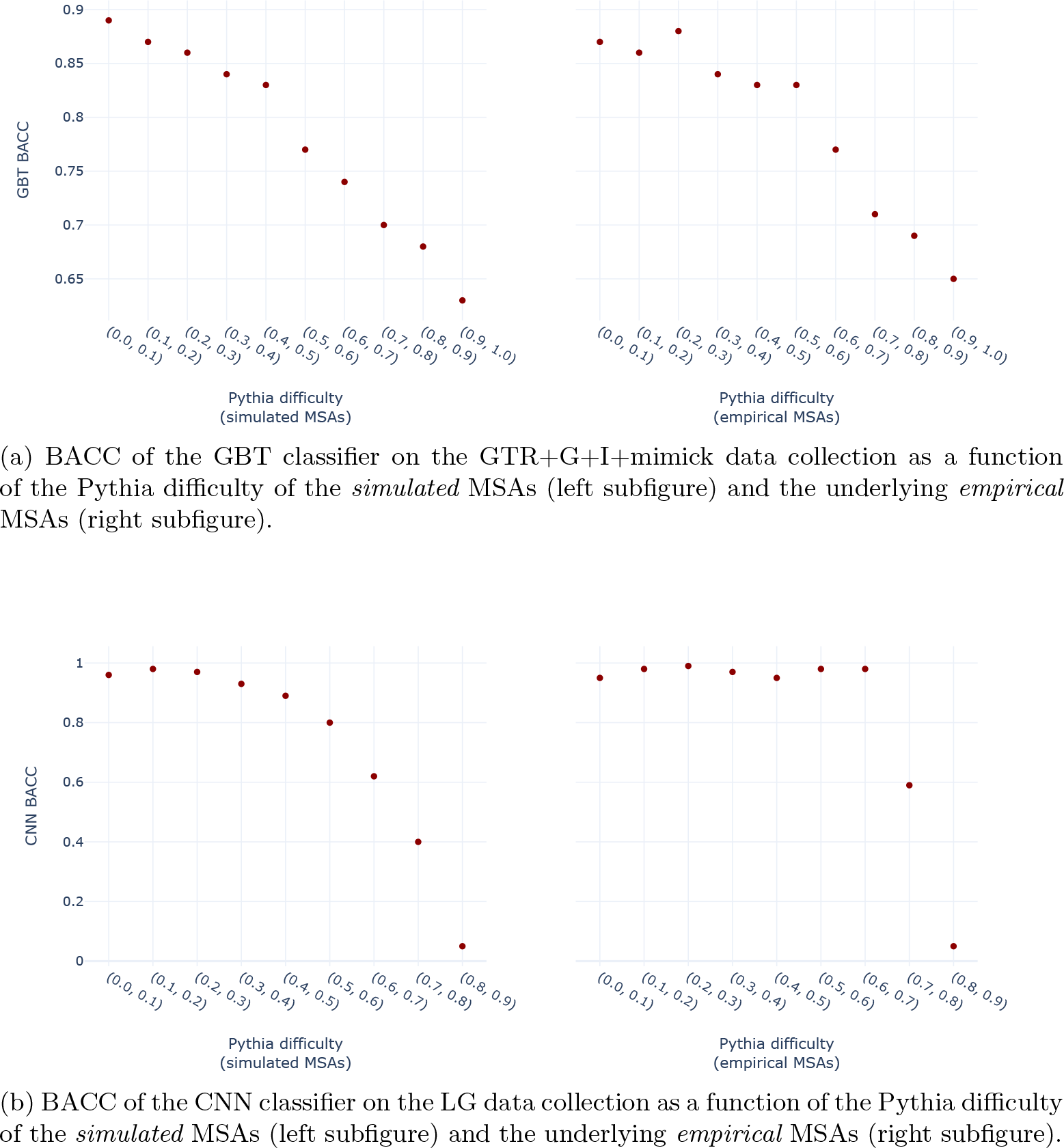
Accuracy of the GBT classifiers depending on the Pythia difficulty of the underlying alignments.

## 4 Discussion and Conclusion

In this study, we assessed the realism of sequence evolution models by attempting to discriminate between simulated MSAs and empirical MSAs using two distinct and independently developed classification methods. Specifically, we evaluated and interpreted the predictive accuracy of these approaches as a measure of realism. By addressing this question, we aimed to gain insights into the ability of current evolutionary models to accurately simulate evolutionary processes using continuous time Markov chains (CTMC). The ability to accurately model sequence evolution and thus simulate realistic MSAs is crucial both for the evaluation of inference tools and the development of neural density estimation techniques for inference.

Note that producing MSAs that are indistinguishable from empirical ones is a necessary but not sufficient condition for the degree of realism of the underlying model. First, poor classification performance can occur because the classifier does simply not deploy appropriate functions or data representations. Hence, one can not guarantee that the simulated MSAs are realistic under all possible criteria. Second, poor performance can also be induced by optimization issues, especially when using deep learning methods. During our experiments, we observed low accuracies for CNNs several times. We managed to alleviate them by adapting the learning rate, the number of filters, or the pooling method, for instance. We thus advise researchers interested in classification performance as a realism metric to closely monitor indicators of poor optimization, in particular, learning curves and gradient norms — in our case, poor optimization also led to a larger variance across folds and discrepancies in accuracy for the two classes. Because we found that all simulated MSAs were easy to discriminate from empirical MSAs, and because our results are consistent across two technically substantially distinct and independent classification methods, we conclude with confidence that the simulated MSAs generated in our study are not realistic.

It is worth noting here that we originally chose to develop a CNN for the classification task, as it is able to capture local dependencies among sites. With a kernel size greater than one, the network could potentially benefit from these dependencies for classification, as they are present in empirical MSAs yet cannot be replicated with standard site-independent models of sequence evolution. However, we discovered that for protein data, the CNN yields accurate performance, even with a kernel size of one in combination with global average pooling (as an alternative to the commonly used local maximum pooling). This type of network primarily focuses on capturing global features while overlooking local among-site dependencies. Consequently, these choices enabled us to thoroughly explore the limitations of current sequence evolution simulation approaches and different evolutionary models beyond their unrealistic assumption of independently evolving sites. However, in the future, a CNN architecture could be deployed to assess the importance of local site dependencies not accounted for in current state-of-the-art simulators.

Our study uses two fundamentally different classifiers, which allows for a broader assessment of possible weaknesses of current sequence evolution simulations: GBTs rely upon diverse, yet welldefined MSA properties, such as branch lengths or the randomness features that take into account the assumption of homogeneity along MSAs in standard simulations. Given the high feature importance of the evolutionary rates (SCC) in the MSA, our GBTs exploit a lack of structure along simulated MSAs. The CNN only considers site-wise composition vectors, and thus exploits a signal that is not directly exploited by the GBTs. Furthermore, for the classification we used diverse and representative empirical protein and DNA databases: TreeBASE comprises representative data sets that are commonly analyzed in the field because it only contains MSAs of published studies, whereas HOGENOM offers a diverse sample of existing data, drawing from 499 nuclear Bacterial genomes, 46 from Archaea, and 121 from Eukaryotes.

The structure detected by our GBTs in empirical nucleotide alignments from TreeBASE is not due to the type of genetic code present. We computed the number of stop codons in all genes in the database and at all three phases, and did not observe an excess of alignments with 0 or 1 stop codons per sequence (Supplementary Figure S17). Instead, it seems to correspond to the pattern found by Bricout et al. [10]. However, in the future it will be interesting to investigate the realism of existing codon models, on a data set of coding DNA sequences.

We used phylogenetic trees reconstructed from these empirical data collections to simulate data as realistically as possible. Thereby, we circumvented having to simulate realistic trees and can invoke simulations that are as similar as possible to the empirical MSAs. However, it is important to note that the realism of the simulations depends on the quality of the inferred phylogenetic trees when deploying this procedure. Since we do not know the true trees of the empirical MSAs, we must acknowledge that there is some uncertainty or error in the inferred trees that the simulations inherit. Hence, at least part of the classifier accuracy, that is, part of the difference between the simulations and the empirical MSAs, could be attributed to the difference between the inferred trees and the true unknown trees. However, our choice to use Maximum Likelihood trees inferred under the same models used for the subsequent simulation (except for the protein data, see below) may constitute the most realistic approach toward generating alignments that resemble empirical MSAs. Indeed, the best-known Maximum Likelihood tree 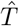 under model M for an alignment A is the best tree we can find that maximizes the probability of observing A. Any other tree is less likely to have generated A under model M (assuming optimization did find the ML tree). Therefore, by simulating with model M along tree 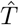, we maximize the probability (or get close to maximizing it) of generating alignment A. We expect that thereby, we also obtain a high probability of generating alignments that resemble A, that is, MSAs that “look” empirical.

However, for protein data, the inference of trees from protein MSAs without indels was performed under the LG substitution model. The resulting trees may be different from the Maximum Likelihood tree obtained under the WAG model or under mixture models. In particular, trees inferred under the LG model may have branches that are too short to be used for simulating MSAs with site-heterogeneous mixture models, because inference with mixture models typically yields longer branches than inference under the LG model. However, looking at amino acid diversity per site (Figure S6) reveals that sites simulated using mixture models look more like empirical sites than sites simulated with LG. Therefore, it remains unclear why mixture models failed to improve alignment realism according to our classifiers. Overall, for some of our experiments on protein data, the mismatch between substitution models used to infer the trees and those employed to simulate the MSAs may be consequential and warrants further investigation.

The classification task was not difficult, neither for DNA nor for protein data. Our CNN achieved an average BACC of 0.98 across all evolutionary models. This shows that existing models of sequence evolution fail to capture important characteristics of empirical site-wise compositions. In turn, this questions to which extent previous results obtained on simulated data apply to empirical data.

We originally hypothesized that with increasing evolutionary model complexity, classification performance would decrease. However, our results do not fully confirm this initial hypothesis. On the contrary, both classifiers remained highly accurate on the most complex evolutionary models for protein simulations with heterogeneous stationary distributions across sites. On DNA simulations, the inclusion of rate heterogeneity and a proportion of invariant sites did not lead to a substantial decrease in CNN classification accuracy, either. Using the HKY substitution model instead of the JC model did also not result in more realistic simulations as a function of observed classification performance. Finally, the most simple models, JC and Poisson, were classified with ease.

We used a state-of-the-art indel model with individual parameters for insertions and deletions and sampled indel parameters from approximated joint distributions. Nevertheless, both classifiers could again easily distinguish simulated from empirical MSAs. In fact, classification accuracy substantially increased on DNA data *with* indels compared to data without indels (GTR+G+I). In contrast, using the *mimick* procedure to superimpose gaps onto simulated data appeared to result in more realistic MSAs. Yet, these MSAs could still be easily identified as simulated ones based on their site-wise compositions, as shown by the CNN results.

Furthermore, the prediction accuracy for protein data tended to be higher than the prediction accuracy for DNA data. We suspect that this is due to the higher number of states in the protein alphabet and therefore the increased number of possible patterns in a protein MSA, which makes it harder to simulate realistic data.

Our findings suggest that existing evolutionary models might not be able to generate data collections that appropriately resemble global low level site composition features of empirical DNA or protein data collections using standard site- and position-independent Continuous Time Markov Chains. Considering the high importance of randomness related features for the GBT classifiers, and the respective feature value distributions, we conclude that the rate of evolution across sites of simulated MSAs are generated more uniformly along the MSA compared to empirical MSAs. For instance, we found that current models cannot reproduce the serial correlation of evolutionary rates that is present in empirical MSAs. We further observe that the proportion of invariant sites in standard simulations reduces their realism as measured by GBT. In addition, the CNN results reveal that simulated alignments have unrealistic properties in terms of site-wise compositions that are independent of correlations among neighboring sites.

The unexpectedly high accuracy of the logistic regression model on simulations under mixture models that produce heterogeneous stationary distributions across sites indicates that these models simulate alignments with an average MSA composition which is distinct from that of empirical data. This is particularly surprising for the LG+S256 models, which had been trained on HOGENOM data [42]. This discrepancy is unlikely to arise from simulating on trees inferred under the LG model rather than mixture models. Indeed shorter branches in the LG trees should result in lower AA diversity per site. However, we did not observe this in our data collections, as sites in simulations under the LG model have slightly higher AA diversity than those in empirical data (Figure S6). Moreover, the site-wise AA diversity appeared similar between simulations under LG+S256 and empirical data. The causes of the discrepancy in average MSA compositions needs to be further investigated.

We believe that in the years to come, learning-based, likelihood-free approaches are likely to be more widely used in our field. Especially, if their performance (both in terms of phylogenetic reconstruction accuracy and runtime) is superior. However, we further believe that likelihood-based inference will continue to play an important role in the area of computational phylogenetics, as the statistical properties of maximum likelihood and MCMC methods for posterior estimation still benefit from a better empirical knowledge.

Looking forward, this work paves the way for approaches to simulate more realistic alignments by developing more realistic models of sequence evolution. We conclude that a substantial amount of research remains to be conducted for improving substitution as well as indel evolution models, for both protein and DNA data.

## 5 Acknowledgements

We thank Philippe Veber, research engineer at the Laboratoire de Biométrie et Biologie Évolutive, for providing us with the source code of his simulator. We gratefully acknowledge support from the CNRS/IN2P3 Computing Center (Lyon - France) for providing computing and data-processing resources needed for this work. This work was granted access to the HPC resources of IDRIS under the allocation 2022-AD011011137R2 made by GENCI. This work was financially supported by the Klaus Tschira Foundation, the ANR grants EVOLUTHON ANR-19-CE45-0010 and PIECES ANR-20-CE45-0017, and by the European Union (EU) under Grant Agreement No 101087081 (Comp-Biodiv-GR).

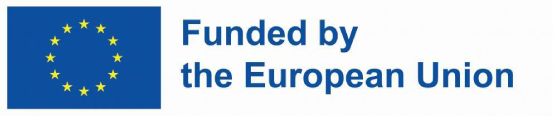

