## Supplementary Material for "Simulations of sequence evolution: how (un)realistic they are and why"

5 **Comparing Evolutionary Models** 24

6 **TreeBASE Data** 25

### 1 Alignment Simulations

Figures S1 and S2 depict schematic overviews of our simulation procedures to obtain the simulated DNA and protein alignments.

### 2 Empirical Parameter Distributions

#### 2.1 Protein Data Collections

##### 2.1.1 Empirical Phylogenetic Trees

We obtained the phylogenies for protein simulations with indels from the HOGENOM database. They were reconstructed by Penel et al. [7] using IQ-TREE [5] under the LG rate matrix, the posterior mean site frequency (PMSF) profile mixture model, and four $\Gamma$ rate categories with 1000 bootstrap replicates.

##### 2.1.2 MSA Length and $\alpha$ Parameter

We obtained MSA lengths and $\alpha$ shape parameters of the $\Gamma$ model of rate heterogeneity from approximated empirical probability density functions (PDFs). The PDF estimate is the linear interpolation of histogram bins of the empirical $\alpha$ parameters and MSA lengths, respectively. We approximated the optimal number of bins automatically using the "auto" method of Harris et al. [3]. For the distribution of $\alpha$ parameters we obtained 21374 bins, with many empty bins due to outliers, and for the MSA lengths distribution we used 38 bins. In Figure S3 we show the distribution of empirical MSA lengths, as well as the distribution of 7000 MSA lengths sampled from the PDF estimate. We show the same for the $\alpha$ parameters, whereas here we removed the 0.99 quantile for better visibility. In both cases, the approximated distribution matches the empirical one very accurately.

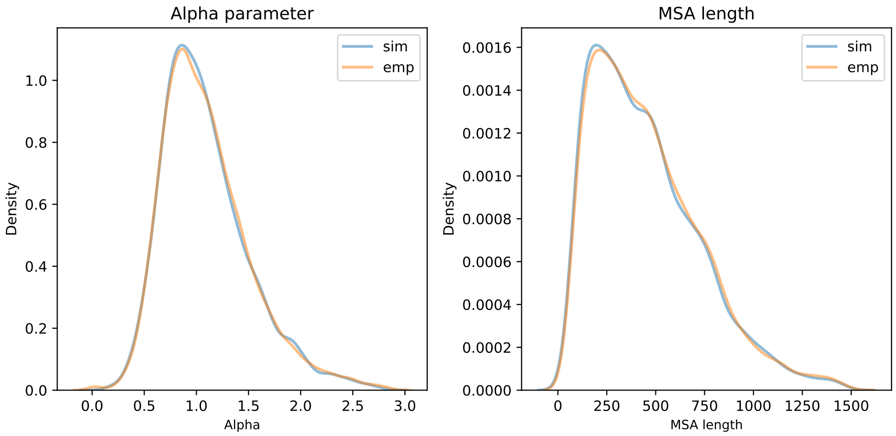

Figure S3: MSA lengths distribution and $\alpha$ parameters distribution for the $\Gamma$ model of rate heterogeneity.

##### 2.1.3 Indel Parameters

We inferred indel parameters from a sample of 652 empirical protein MSAs with more than 20 sequence using SpartaABC, they are referred to as empirical parameters in this section. We then deduced the joint PDF from these parameters via kernel density estimation (KDE) using a bandwidth of 0.4867 as explained in the paper in Section 2.1. To simulate MSAs, we drew parameters from the approximated PDF. They are referred to as simulated parameters. Here, the approximated distribution is represented by the density function based on a sample of 7000 values for each parameter type. To compare empirical and simulated parameters, we additionally show the ECDF of each parameter type for both. The ECDF shows the proportion of occurrences of each unique value in the dataset. In contrast to a histogram or density plot, it does not require a binning or smoothing parameters. In this case this is especially useful as the sample size of empirical parameters is small.
The comparison of ECDFs in Figure S4 indicates a good match between empirical and simulated distributions, except for the deletion rate parameter which is overestimated by the approximated PDF, which is most evident when examining percentiles within the 0.1 to 0.8 range.

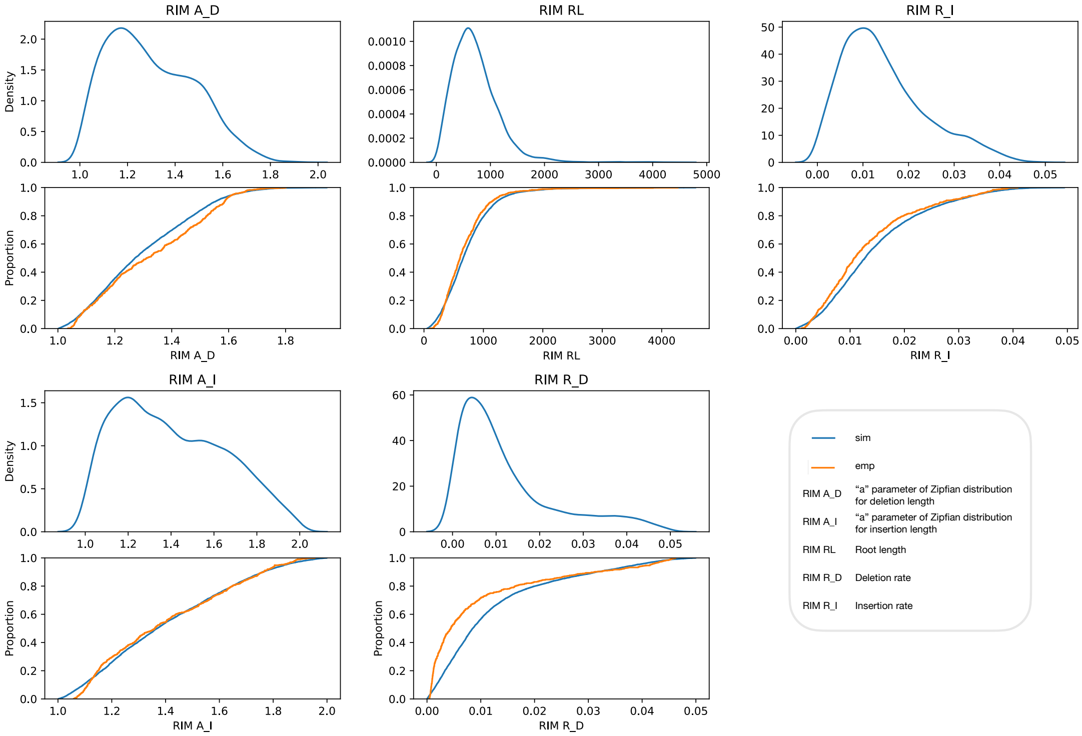

Figure S4: Protein indel parameter distributions represented by the density function (top) and ECDF (bottom) for each parameter type. Parameters are drawn from the approximated PDF (sim) based on inferred parameters from empirical MSAs (emp).

In addition to the inferred Indel parameters, we also analyzed the resulting MSA lengths from data simulated based on the inferred PDF, as compared to MSA lengths from the empirical data collection. We constrained the simulated MSAs to contain a minimum of 100 and a maximum of 10,000 sites to match the MSA length range of our empirical data after removing outliers. To do so, we repeated the simulation if the resulting MSA length exceeded these limits. To validate this approach, we show the density functions of MSA lengths for both empirical and simulated data in Figure S5, where the simulated data is generated using the $LG + S256 + GC + sparta$ model.

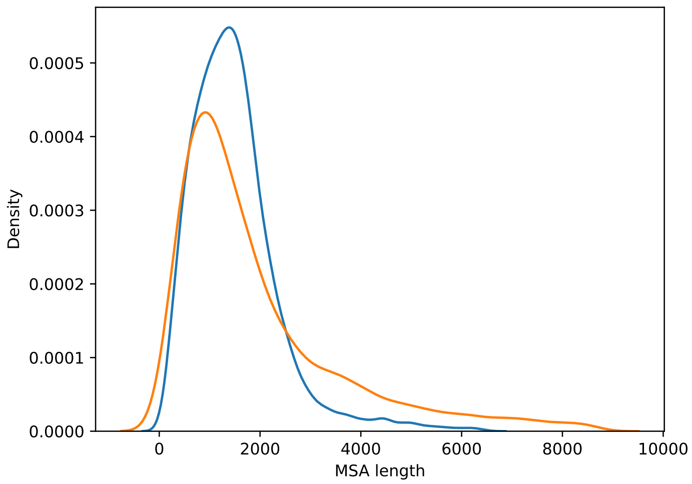

Figure S5: Density function of MSA lengths of the empirical data collection with Indels (emp) and of the simulated protein data collection under the $LG + S256 + GC + sparta$ model (sim).

In Figure S5 we observe an underestimation of the average MSA length in the simulated data, which might be attributed to the overestimated deletion rate as observed in Figure S4. Additionally, the simulated distribution has a wider and longer tail than the empirical distribution, which could be due to mismatching indel-rate-ratios when drawing from the joint distribution.

###### 2.1.4 Site-wise Amino Acid diversity

Here we investigated the number of unique AAs per site, within the empirical HOGENOM data collection without indel and simulations under LF and LG+S256 (a mixture model with 256 profiles). For this, we sampled 100,000 sites across all sites of all MSAs for each data collection and counted the number of unique AAs in the sampled sites.

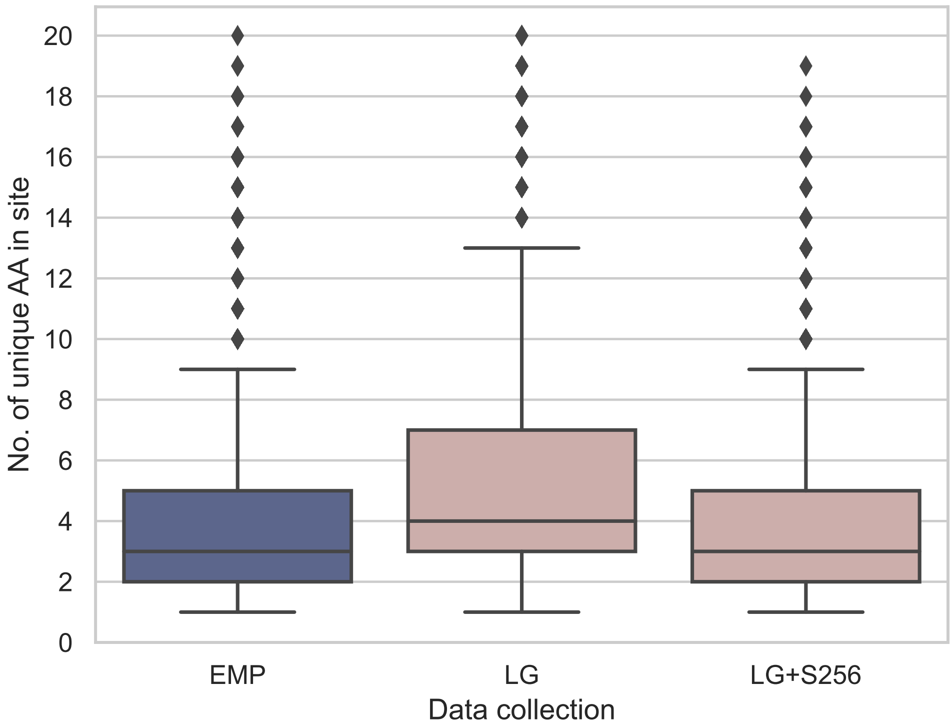

Figure S6: Site-wise AA diversity as the number of unique AAs per site.

#### 2.2 DNA Data Collections

##### 2.2.1 Indel Parameters

We used SpartaABC to infer indel parameters from a sample of 5060 DNA MSAs from TreeBASE with a number of sequences above 20. Similarly to the descriptions in Section 2.1.3, we estimated the PDF from the inferred empirical parameters, drew 7000 simulated parameters, and plotted the ECDFs for every parameter type (see Figure S7). Again, we can observe a good match between estimated empirical and simulated distributions.

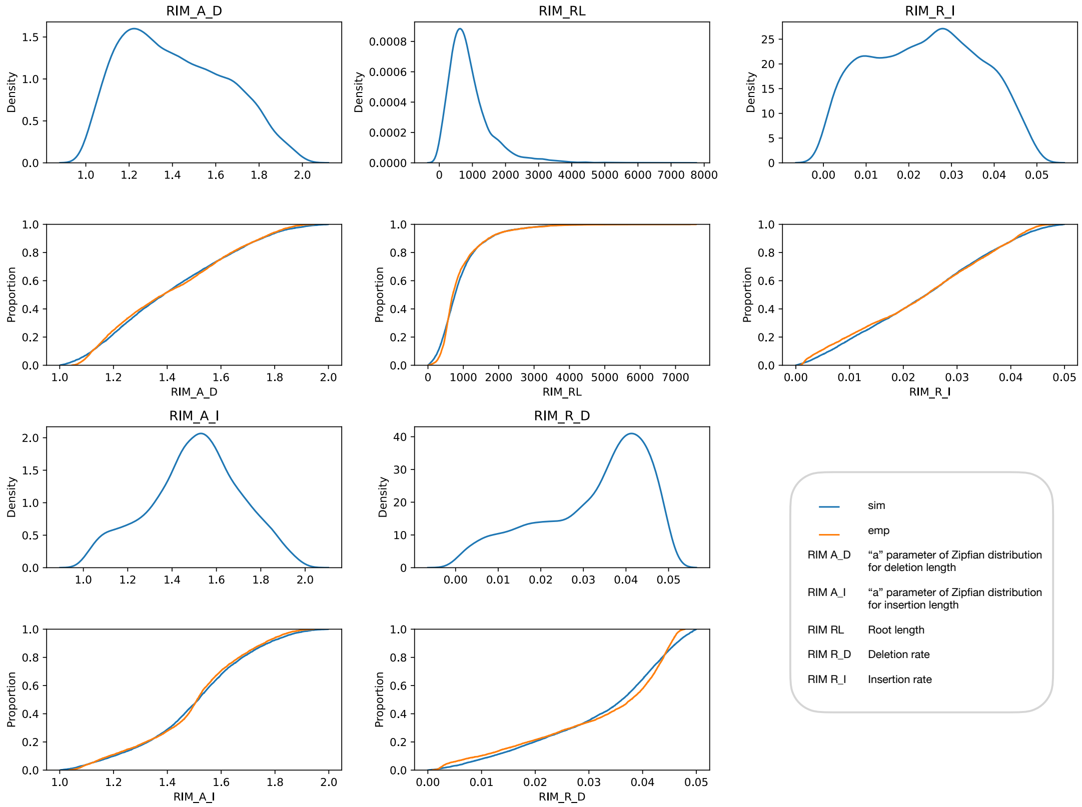

Figure S7: DNA indel parameter distributions represented by the density function (top) and ECDF (bottom) for each parameter type. Parameters are drawn from the approximated PDF (sim) based on inferred parameters from empirical MSAs (emp).

Further, we examined the PDFs of empirical and simulated MSA lengths resulting from the simulations of DNA MSAs under the *GTR+G+I+sparta* model (see Figure S8). In comparison to the simulated protein MSAs, here, the simulated MSAs are shorter on average than the empirical MSAs.

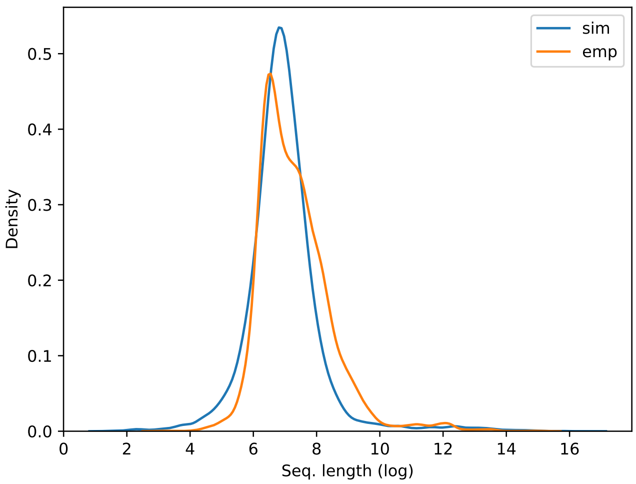

Figure S8: Density function of MSA lengths of the empirical data collection with Indels (emp) and of the simulated DNA data collection under the *GTR+G+I+sparta* model (sim).

### 3 CNN Training and Performance

All aspects of our classifier, including data preprocessing, model architecture, as well as training and evaluation were implemented in Python [10]. We employed PyTorch [6], a popular open-source machine learning framework, to implement our model and trained it on an Nvidia Tesla V100 GPU.

##### 3.0.1 Parameter optimization

We optimized the learning rate and the number of filters. For the former, we used the learning rate range test (LRRT) [9] on learning rates from $10^{-7}$ to 0.5 for each data collection and fold independently. Based on the validation loss after 100 epochs and the learning curves, we then chose a learning rate. Through this procedure, we determined an optimal learning rate of 0.001 for all DNA data collections. In the case of protein data, we determined varying learning rates ranging from 0.001 to 0.0117 across different data collections. Here, the optimized learning rates correspond to one-tenth of the upper bound identified by the LRRT, which proved to be the most effective choice for the protein data collections. Exact learning rates are reported in the configuration files of the trained models, which are available at https://github.com/JohannaTrost/seqsharp. We used a batch size of 128 for protein data and 64 for DNA data.

#### 3.1 Network Evaluation

In this section, we show the learning curves for all Convolutional Neural Networks (CNNs) used in our study, which are divided into two groups: those trained on DNA data (Figure S9) and those trained on protein data (Figure S10). For each evolutionary model used in the simulations, we show both the loss and balanced accuracy score (BACC) over epochs on both validation and training data, with all 10 folds stacked together. Note that the training of different folds of the cross-validation procedure might stop at different epochs, as each fold is trained individually.

For the DNA data, we observed stronger fluctuations during the training process, as well as more variation between folds compared to the protein data. Moreover, the DNA-networks tend to converge more slowly, when simulations were based on the GTR substitution model (Figure S9). Among the CNNs for DNA data collections, we found that learning curves for simulations under the JC and HKY models reached a plateau earliest and exhibited the least variation across folds.

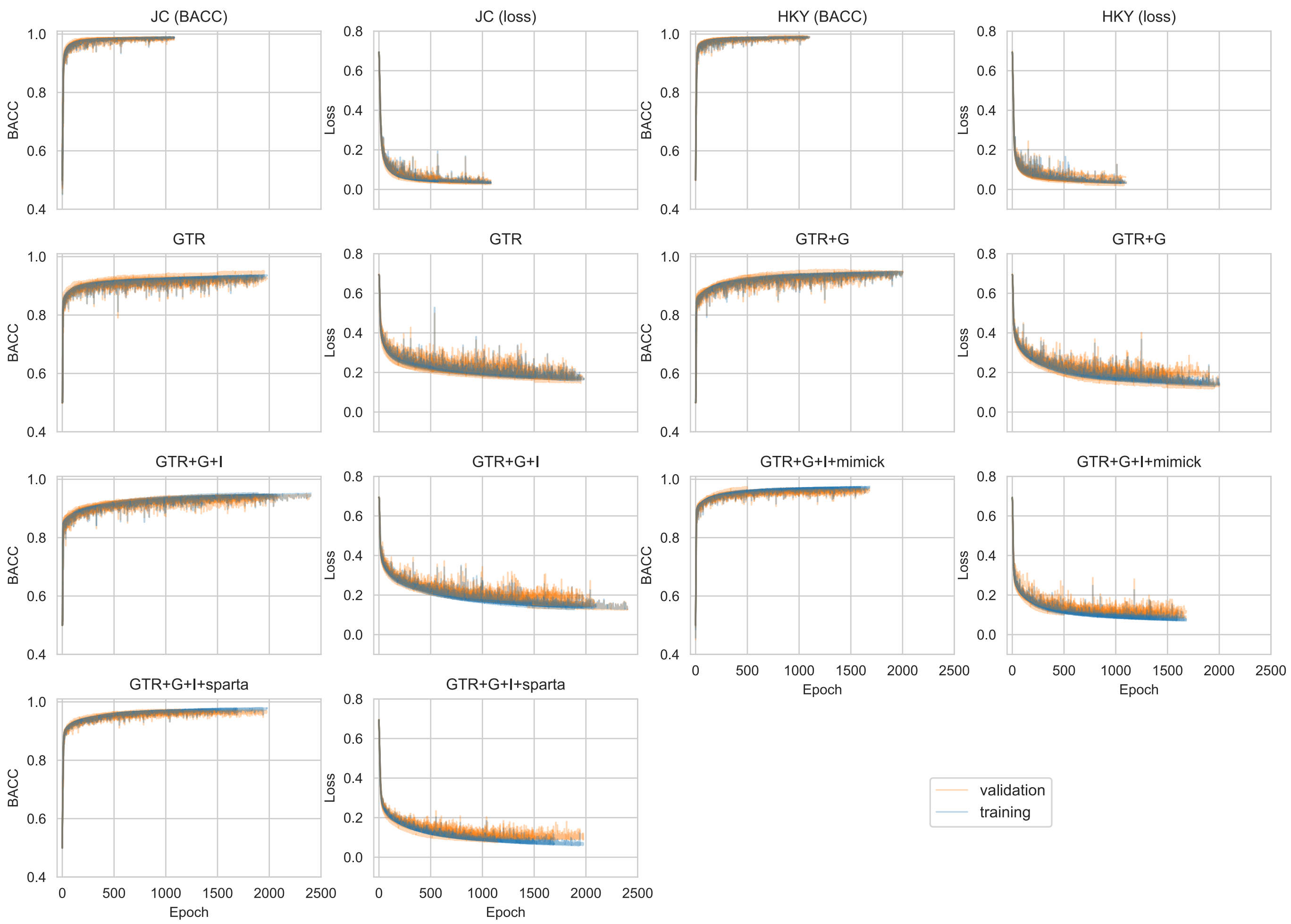

Figure S9: Learning curves of CNNs trained on DNA data collections.

When training CNNs on protein data collections, we observed a general trend of a rapid improvement in performance within the first 100 epochs, followed by only minor gains thereafter. Interestingly, we found that models trained on simulations under the *S256* and *C60* models, which exhibit heterogeneity in stationary distributions across sites, had very similar learning curves with little variation across folds. In contrast, apart from the Poisson model, we observed more variation in learning curves for CNNs trained on site homogeneous models, that are *LG* and *WAG*. Moreover, the validation and training curves were closely aligned for all simulation setups, indicating a robust performance.

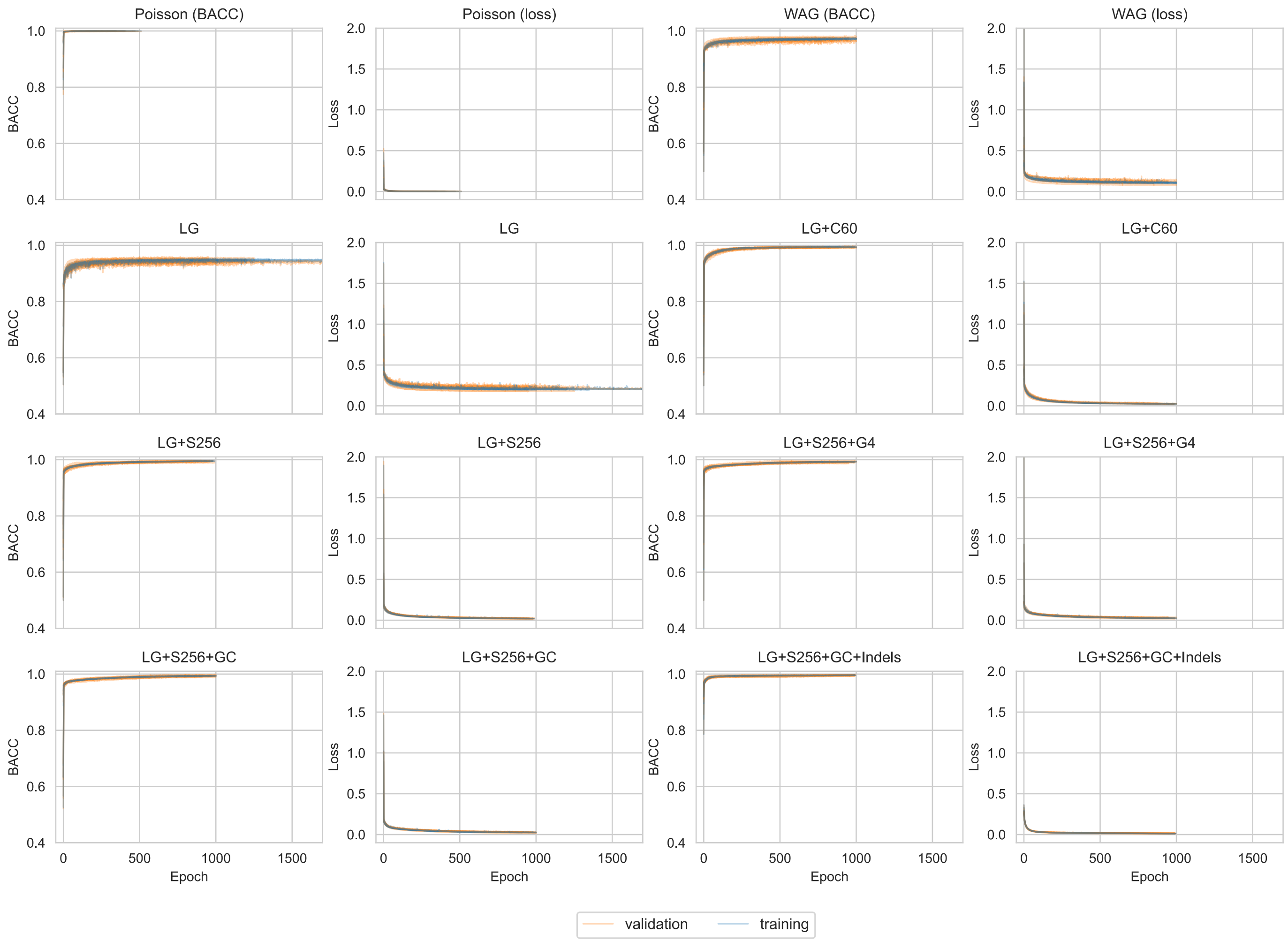

Figure S10: Learning curves of CNNs trained on protein data collections.

It should be noted that, surprisingly, the performance for the validation data is slightly better than for the training data, in at least one fold for all types of simulations. In particular, this phenomenon was observed in three folds for simulations with the Poisson, LG+S256, and JC models. For the other CNNs, this phenomenon was observed in only one to two folds. This effect occurred mainly in three specific splits (folds) across the different simulation configurations.

Furthermore, in some instances, the training process was unexpectedly interrupted and required resumption, such as in the case of the CNN model trained on the GTR model simulations at epoch 1500. Upon resuming training, we noticed a slight fluctuation in the learning curve, which is barely noticeable in the graphs. Despite our efforts, we were unable to determine the cause of this sudden leap. However, we believe it had little impact on the final average BACC that we report.

In addition to the learning curves, we considered multiple measures to validate the trained models and analyze its performance (see Table S1). Here, solely the performance on validation data was taken into account.
Apart from the average BACC across folds (*BACC*) we considered the standard error of the BACC (*SE*), which is an estimate of the true dispersion across folds. The BACC is more dispersed across folds as the average BACC decreases: for a BACC greater than 0.98 the SE is less than 0.001, whereas for a BACC less than 0.98 the SE is greater than 0.001. This suggests that as the BACC increases, the CNN also generalizes better, since the BACC becomes more consistent across folds.

To ensure that the network is not overfitted at the epoch with the maximum validation BACC, we also take into account twelve epochs surrounding the selected one (*n-BACC*) averaged over folds. If the selected epoch is e.g. the last one, we only consider the six preceding epochs. We report the maximum, minimum, and mean across these neighboring BACCs. Overall, we find that the difference between the neighboring BACCs and the best BACC does not indicate an over-fit at the selected epoch. We further report the accuracy per class (*ACC SIM* and *ACC EMP*) and their mean absolute error across folds (*Class MAE*) to evaluate the consistency of the model's performance across the two classes. We found a class MAE of less than 0.01 in most cases. Therefore, the BACC can be used as a metric for evaluating the realism of simulated data collections so that performance on empirical data is not ignored, but without being distorted by the inaccuracies in either of the two class accuracies.

| Evo. Model | BACC | SE | AVG n-BACC | MIN n-BACC | MAX n-BACC | ACC SIM | ACC EMP | Class MAE |
|---|---|---|---|---|---|---|---|---|
| DNA data | | | | | | | | |
| JC | 0.9873 | 0.0007 | 0.9841 | 0.9835 | 0.9851 | 0.9906 | 0.9841 | 0.0067 |
| HKY | 0.9907 | 0.0007 | 0.9878 | 0.9868 | 0.9888 | 0.9906 | 0.9908 | 0.0051 |
| GTR | 0.9303 | 0.0027 | 0.9235 | 0.9208 | 0.9260 | 0.9438 | 0.9168 | 0.0270 |
| GTR+G | 0.9427 | 0.0028 | 0.9348 | 0.9312 | 0.9369 | 0.9500 | 0.9353 | 0.0180 |
| GTR+G+I | 0.9420 | 0.0022 | 0.9343 | 0.9327 | 0.9361 | 0.9494 | 0.9345 | 0.0202 |
| mimick | 0.9686 | 0.0010 | 0.9625 | 0.9602 | 0.9641 | 0.9694 | 0.9677 | 0.0070 |
| sparta | 0.9703 | 0.0015 | 0.9637 | 0.9602 | 0.9661 | 0.9761 | 0.9645 | 0.0120 |
| Protein data | | | | | | | | |
| Poisson | 0.9996 | 0.0002 | 0.9990 | 0.9987 | 0.9994 | 0.9999 | 0.9994 | 0.0007 |
| WAG | 0.9736 | 0.0016 | 0.9699 | 0.9692 | 0.9710 | 0.9753 | 0.9719 | 0.0121 |
| LG | 0.9507 | 0.0018 | 0.9439 | 0.9420 | 0.9448 | 0.9522 | 0.9491 | 0.0152 |
| LG+C60 | 0.9942 | 0.0005 | 0.9922 | 0.9917 | 0.9925 | 0.9954 | 0.9930 | 0.0036 |
| LG+S256 | 0.9952 | 0.0006 | 0.9938 | 0.9935 | 0.9940 | 0.9961 | 0.9943 | 0.0024 |
| LG+S256+G4 | 0.9930 | 0.0007 | 0.9914 | 0.9911 | 0.9917 | 0.9937 | 0.9924 | 0.0039 |
| LG+S256+GC | 0.9938 | 0.0008 | 0.9925 | 0.9920 | 0.9932 | 0.9968 | 0.9908 | 0.0063 |
| sparta | 0.9959 | 0.0004 | 0.9946 | 0.9945 | 0.9949 | 0.9973 | 0.9945 | 0.0036 |

Table S1: Additional measures to validate the CNN performances on simulations under the different evolutionary models. Parameter configurations of simulations (*Evo. Model*) are sorted with increasing complexity from top to bottom for both DNA and protein data. For both, the last row shows results on data collections with Indels: *mimick* refers to the model GTR+G+I+mimick and sparta to GTR+G+I+sparta and LG+S256+GC+sparta respectively. *BACC* is the average BACC and *SE* the standard error across folds. *n-BACC* denotes the BACCs of the neighboring epochs of the selected (i.e. best) epoch.

As stated in the main paper, we excluded the first two sites of the empirical protein data collections to prevent biasing the prediction. Although the first site of protein sequences typically contains a Methionine (M), which is absent in simulated protein sequences, there is little evidence to suggest that the second site could also introduce bias. However, since all models were trained on data with the second site removed, we conducted a sanity check to ensure that its exclusion did not significantly affect our resulting BACC. Specifically, we tested the CNN pretrained on empirical data with the second site excluded against simulations under the LG and LG+S256 models on empirical data with and without the second site. The results of these tests are presented in Table S2, where we observe that the empirical accuracies of the two tests differ only in the third and fourth decimal place.

| Evo. Model | EMP ACC | EMP ACC | ABS DIFF |
|---|---|---|---|
| | Without 2nd site | With 2nd site | |
| LG | 0.92828 | 0.92784 | 0.00043 |
| LG+S256 | 0.99354 | 0.99326 | 0.00029 |

Table S2: Comparison of the accuracy on empirical protein data with and without the second site.

### 4 Gradient Boosted Trees

#### 4.1 Feature Computation and Training Procedure

We generated the training data and features using a custom pipeline that we programmed using Snakemake [4] and Python 3. For each MSA, we performed a single RAxML-NG tree inference based on a random starting tree using RAxML-NG's `--search1` execution mode. On the resulting maximum likelihood tree, we executed the RAxML-NG `--eval` mode. In this run mode, RAxML-NG does not alter the tree topology and only optimizes the branch lengths and substitution model parameters. Based on the resulting maximum likelihood tree, we compute the branch length features $brlen_{min}$, $brlen_{max}$, $brlen_{avg}$, $brlen_{std}$, $brlen_{med}$, and $brlen_{sum}$. We used PyPythia [2] to compute the following set of features: *number of taxa*, *number of sites*, *number of patterns*, *sites-over-taxa ratio*, *patterns-over-taxa ratio*, *% invariant*, *Entropy*, *Bollback multinomial*, *Pattern entropy*, *difficulty*, *parsimony RF-Distance*, and *% parsimony unique* (see below for further details on the computation of the *Entropy*, *Bollback multinomial*, and the *Pattern entropy*). Note that we do not use the number of taxa and the number of sites as prediction features. Based on these features, we additionally compute the *patterns-over-sites ratio* as the number of patterns divided by the number of sites. To compute the randomness features, we used the Fourmilab Random Sequence Tester (FRST) (https://www.fourmilab.ch/random/).

In our setup, we used RAxML-NG version 1.1.0 and PyPythia version 1.0.0. The pipeline code and instructions to reproduce the results, as well as Apache parquet files containing all training data, are available in our GitHub repository at https://github.com/tschuelia/SimulationStudy.

We executed the pipeline on two servers at the Heidelberg Institute of Theoretical Studies. The two servers have the following hardware specifications: Xeon Platinum 8260, 48 cores with 2.4 GHz and 754 GB RAM; AMD EPYC 7413, 24 cores with 2.65 GHz and 512 GB RAM.

We trained all LightGBM (LGB) GBT classifiers using a custom training script that is also available at the linked GitHub repository, alongside all training results presented in this paper. We executed the training script on an Apple MacBook with M1 Pro Chip and 16 GB RAM. We used 100 iterations of Optuna to optimize hyperparameters of the classifiers. Table S3 shows the initial setting as well as the tested value range for all optimized parameters. We set all other parameters to the default setting in LGB version 3.3.2.

| LGB hyperparameter identifier | initial setting | tested value range |
|---|---|---|
| learning rate | 0.05 | $[0.01, 0.25]$ |
| max_depth | 10 | $[5, 10]$ |
| lambda_l1 | $10^{-6}$ | $[10^{-8}, 10.0]$ |
| lambda_l2 | 0.1 | $[10^{-8}, 10.0$ |
| num_leaves | 20 | $[2, 20]$ |
| bagging_fraction | 0.9 | $[0.4, 1.0]$ |
| bagging_freq | 4 | $[1, 7]$ |
| min_child_samples | 50 | $[30, 100]$ |

Table S3: Parameter settings for the GBT hyperparameter optimization using the Optuna framework.

**Entropy** We define the Entropy $H$ of an MSA as the average Shannon Entropy over all sites of the MSA. Let $N$ be the number of taxa in the MSA and $M$ the number of sites. The Entropy then is computed as

$$H(\text{MSA}) = \frac{1}{M} \sum_{i=1}^{M} H(\text{site}_i) \tag{1}$$

$$\text{with} \quad H(\text{site}_i) = -\sum_{j=1}^{N} P(\text{char}_{j,i}) \cdot log(P(\text{char}_{j,i})) \tag{2}$$

where $P(\text{char}_{j,i}$ is the frequency of character $j$ at site $i$.

**Bollback multinomial** The Bollback multinomial is a multinomial test statistic that quantifies the frequency of patterns in an MSA. In our setup, we use the test statistic as defined by Equation 7 in [1]:

$$T(\text{MSA}) = \left( \sum_{i=1}^{n} N_{\xi(i)} \cdot ln(N_{\xi(i)}) \right) - M \cdot ln(M) \tag{3}$$

Where $M$ denotes the number of sites in the MSA, $n$ the number of unique site patterns, $\xi(i)$ the $i$-th unique pattern, and $N_{\xi(i)}$ the number of occurrences of pattern $\xi(i)$.

**Pattern entropy** The summation in Equation (3) is similar to the computation of the Shannon entropy. Since the *Bollback multinomial* subtracts the number of sites of this entropy-like metric, and thus might be biased by the length of the MSA, we decided to add the entropy-like summation of pattern frequencies as a separate feature. Thus, we compute the *Pattern entropy* as

$$T(\text{MSA}) = \sum_{i=1}^{n} N_{\xi(i)} \cdot ln(N_{\xi(i)}) \tag{4}$$

Where $M$ denotes the number of sites in the MSA, $n$ the number of unique site patterns, $\xi(i)$ the $i$-th unique pattern, and $N_{\xi(i)}$ the number of occurrences of pattern $\xi(i)$.

##### 4.2 Monte Carlo Value for Pi

The authors of the Fourmilab Random Sequence Tester (https://www.fourmilab.ch/random/) split every six bytes of the analyzed sequence into 24-bit long $X$ and $Y$ point coordinates inside a $2^{24} \times 2^{24}$ pixel square. If the point $(X, Y)$ is located inside the circle enclosed by the square, the point is considered a *hit*. The proportion of hits to the overall number of points is then used to estimate the value of $\pi$. For long byte sequences, this estimated value converges to $\pi$ if the byte values in the sequence were chosen randomly.

##### 4.3 Substitution Rate Visualization

In Figure S11 we show additional examples of the substitution rate patterns for gapless TreeBASE DNA MSAs. We present substitution rates of empirical MSAs, and MSAs simulated under the GTR model for MSAs with IDs 40-44. The complete set of figures can be accessed at https://github.com/tschuelia/SimulationStudy/tree/master/figures/sub_rates_vis_gapless_dna_gtr.

##### 4.4 Performance and Feature Importance

Table S4 shows the prediction accuracy as well as the three most important features for the gradient boosted tree classifiers using all features presented in the main paper. The features are sorted by importance, meaning the most important feature is listed first. We computed the gain-based feature importance. The gain-based feature importance directly measures the contribution of a feature to the reduction of the loss function.

| Data collection | BACC | Top three important features |
|---|---|---|
| | DNA data | |
| JC | 0.95 | SCC, $brlen_{stdev}$, Entropy |
| HKY | 0.95 | SCC, $brlen_{stdev}$, Entropy |
| GTR | 0.94 | SCC, $brlen_{stdev}$, Entropy |
| GTR+G | 0.65 | SCC, Pattern entropy, $brlen_{sum}$ |
| GTR+G+I | 0.62 | SCC, Pattern entropy, $brlen_{sum}$ |
| GTR+G+I+mimick | 0.70 | SCC, % invariant, patterns-over-sites ratio |
| GTR+G+I+sparta | 0.94 | SCC, % gaps, $mean_{rand}$ |
| | Protein data | |
| Poisson | 0.99 | SCC, $brlen_{med}$, Bollback multinomial |
| WAG | 0.98 | SCC, $brlen_{med}$, % invariant |
| LG | 0.99 | SCC, $brlen_{med}$, % invariant |
| LG+C60 | 0.97 | SCC, Entropy, patterns-over-site-ratio |
| LG+S256 | 0.99 | SCC, Entropy, % invariant |
| LG+S256+G4 | 0.95 | SCC, Entropy, patterns-over-site-ratio |
| LG+S256+GC | 0.95 | SCC, Entropy, patterns-over-site-ratio |
| LG+S256+GC+sparta | 0.98 | patterns-over-site-ratio, SCC, Entropy |

Table S4: Average of the BACC on empirical and simulated data collections across 10 folds. Parameter configurations of simulations listed in the first column are sorted with increasing complexity from top to bottom for both DNA and protein data. The third column lists the three most important features for classifying the data.

Table S5 shows the BACC for all classifiers, alongside the three most important prediction features when training the classifiers using only the MSA based features (i.e. without the branch length features).

| Data collection | BACC (all) | BACC (no brlens) | Top three important features (no brlens) |
|---|---|---|---|
| | | DNA data | |
| JC | 0.96 | 0.96 | SCC, % invariant, $\text{mean}_{\text{rand}}$ |
| HKY | 0.96 | 0.95 | SCC, % invariant, $\text{mean}_{\text{rand}}$ |
| GTR | 0.94 | 0.94 | SCC, % invariant, $\text{mean}_{\text{rand}}$ |
| GTR+G | 0.89 | 0.88 | SCC, Pattern entropy, Entropy |
| GTR+G+I | 0.89 | 0.88 | SCC, Pattern entropy, Bollback multinomial |
| GTR+G+I + mimick | 0.77 | 0.74 | SCC, parsimony RF-Distance, % invariant |
| GTR+G+I + sparta | 0.94 | 0.94 | SCC, % gaps, $\text{mean}_{\text{rand}}$ |
| | | Protein data | |
| Poisson | 0.99 | 0.99 | SCC, % invariant, $\text{Entropy}_{\text{rand}}$ |
| WAG | 0.99 | 0.99 | SCC, % invariant, $\text{Entropy}_{\text{rand}}$ |
| LG | 0.99 | 0.98 | SCC, % invariant, $\text{Entropy}_{\text{rand}}$ |
| LG+C60 | 0.98 | 0.97 | SCC, Entropy, % invariant |
| LG+S256 | 0.99 | 0.98 | SCC, Entropy, $\text{Entropy}_{\text{rand}}$ |
| LG+S256+G4 | 0.99 | 0.97 | SCC, Entropy, % gaps |
| LG+S256+GC | 0.98 | 0.96 | SCC, Entropy, % gaps |
| LG+S256+GC +sparta | 0.99 | 0.98 | % invariant, SCC, % gaps |

Table S5: Average of the BACC on empirical and simulated data collections across 10 folds when using all features (2nd column) and when using only the MSA based prediction features (3rd column), meaning we do not include branch length features in either training or prediction. Parameter configurations of simulations listed in the first column are sorted with increasing complexity from top to bottom for both DNA and protein data. The third column lists the three most important features for classifying the data.

Table S6 shows the BACC for all classifiers, alongside the three most important prediction features when training the classifiers using no randomness features. This table shows that the randomness features are especially valuable for more complex DNA models (GTR+G and GTR+G+I).

| Data collection | BACC (all) | BACC (no randomness) | Top three important features (no randomness) |
|---|---|---|---|
| | | DNA data | |
| JC | 0.96 | 0.95 | $brlen_{sum}$, % invariant, $brlen_{max}$ |
| HKY | 0.96 | 0.94 | $brlen_{sum}$, % invariant, $brlen_{max}$ |
| GTR | 0.94 | 0.94 | $brlen_{sum}$, % invariant, $brlen_{max}$ |
| GTR+G | 0.89 | 0.65 | parsimony RF-Distance, $brlen_{med}$, patterns-over-site-ratio |
| GTR+G+I | 0.89 | 0.61 | parsimony RF-Distance, $brlen_{med}$, % invariant |
| GTR+G+I + mimick | 0.77 | 0.70 | parsimony RF-Distance, patterns-over-site-ratio, % invariant |
| GTR+G+I + sparta | 0.94 | 0.84 | % invariant, Pattern entropy, % gaps |
| | | Protein data | |
| Poisson | 0.99 | 0.99 | % invariant, $brlen_{stdev}$, patterns-over-site-ratio |
| WAG | 0.99 | 0.98 | % invariant, $brlen_{sum}$, parsimony RF-Distance |
| LG | 0.99 | 0.99 | % invariant, $brlen_{sum}$, parsimony RF-Distance |
| LG+C60 | 0.98 | 0.98 | $brlen_{max}$, $brlen_{sum}$, % invariant |
| LG+S256 | 0.99 | 0.99 | $brlen_{max}$, % invariant, $brlen_{sum}$ |
| LG+S256+G4 | 0.99 | 0.96 | $brlen_{max}$, $brlen_{stdev}$, % invariant |
| LG+S256+GC | 0.98 | 0.95 | $brlen_{max}$, $brlen_{stdev}$, % invariant |
| LG+S256+GC +sparta | 0.99 | 0.98 | $brlen_{stdev}$, $brlen_{max}$, % invariant |

Table S6: Average of the BACC on empirical and simulated data collections across 10 folds when using all features (2nd column) and when not using the five randomness features. Parameter configurations of simulations listed in the first column are sorted with increasing complexity from top to bottom for both DNA and protein data. The third column lists the three most important features for classifying the data.

##### 4.5 Substitution Distributions over DNA MSAs

In Figures S12 to S14, we visualize the substitution numbers in DNA MSAs. As written in the main text: We split MSAs site-wise into 100 parts (buckets), averaged the substitution numbers per bucket (normalized with the maximum number of substitutions per MSA), and averaged the buckets over every MSA. We interpolated the bucket averages for MSAs with a number of sites below 100.

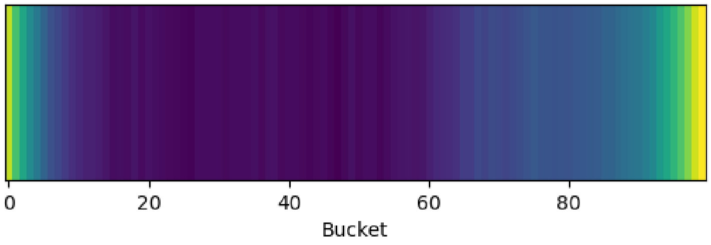

Figure S12: Visualized substitution rates averaged over all empirical DNA MSAs.

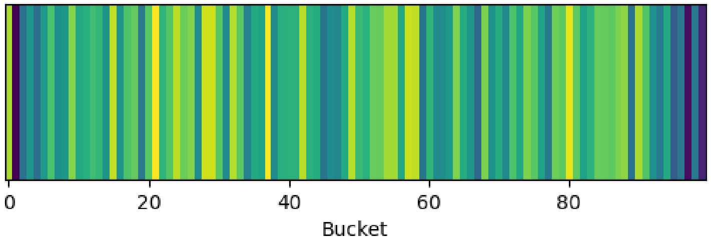

Figure S13: Visualized substitution rates averaged over all DNA MSAs simulated under the model GTR+G+I+sparta.

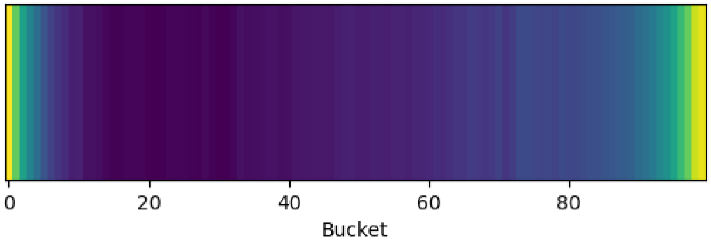

Figure S14: Visualized substitution rates averaged over all DNA MSAs simulated under the model GTR+G+I+mimick.

#### 4.6 Feature Distributions for GTR+G and GTR+G+I

In the main paper, we depicted the distribution of feature values for the proportion invariant, parsimony RF-Distance, as well as the maximum branch length for the JC data collection. While the described trend holds true for all simulated data collections, the pattern is less pronounced the more complex the model of evolution is. Especially for the simulated DNA data under GTR+G and GTR+G+I, the distribution of feature values matches the empirical distribution closely as depicted in Figures S15 and S16.

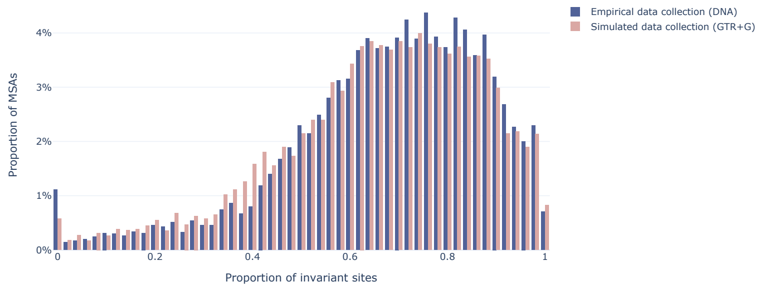

(a) Distribution of proportion of invariant feature values for the GTR+G and empirical data collections.

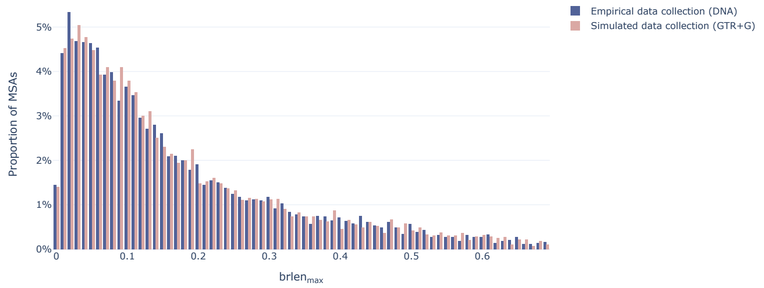

(b) Distribution of *brlen*$_{max}$ feature values for the GTR+G and empirical data collections.

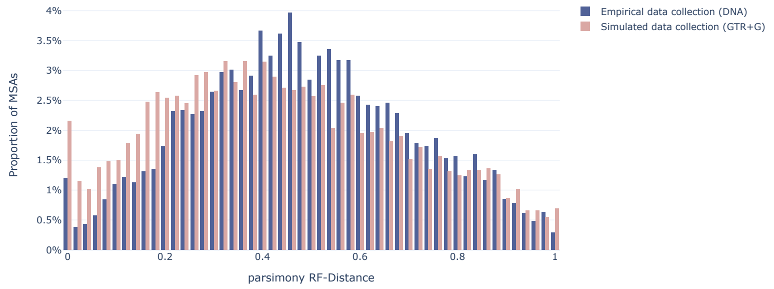

(c) Distribution of *parsimony RF-Distance* feature values for the GTR+G and empirical data collections.

Figure S15: Feature distribution for important features for classifying the GTR+G data collection. The dark blue bars represent the empirical data collection and the light pink bars represent the simulated GTR+G data collection.

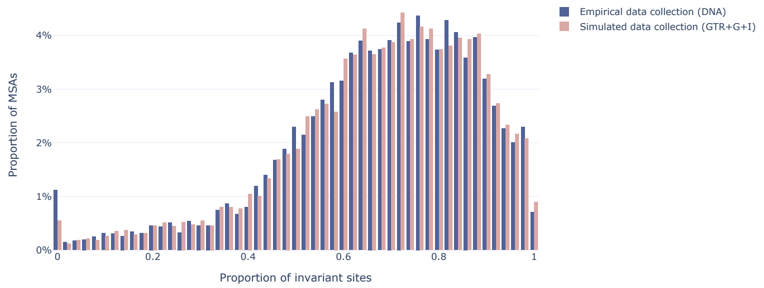

(a) Distribution of proportion of invariant feature values for the GTR+G+I and empirical data collections.

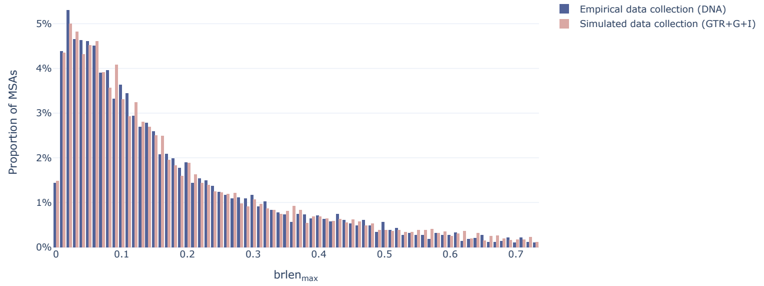

(b) Distribution of *$brlen_{max}$* feature values for the GTR+G+I and empirical data collections.

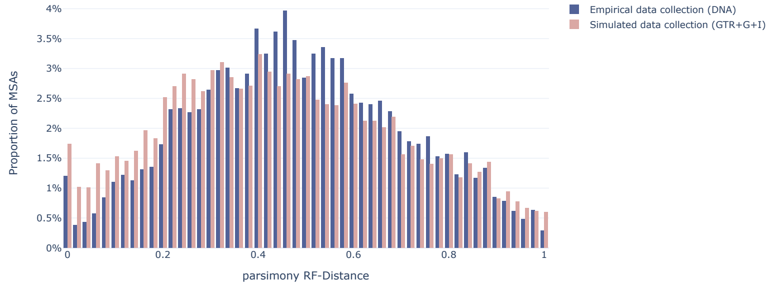

(c) Distribution of *parsimony RF-Distance* feature values for the GTR+G+I and empirical data collections.

Figure S16: Feature distribution for important features for classifying the GTR+G+I data collection. The dark blue bars represent the empirical data collection and the light pink bars represent the simulated GTR+G+I data collection.

### 5 Comparing Evolutionary Models

To compare the BACCs from two different simulated data collections, i.e. evolutionary models, we conducted multiple unpaired two-samples t-tests for DNA and protein networks separately, where one sample consists of the validation BACC for each fold of the respective GBT/CNN. Note that we apply the t-tests for GBT and CNN separately. The null hypothesis is that the BACCs are equal. We use the standard significance cutoff $\alpha = 0.05$. We discard the null hypothesis if the respective P-value is $< \alpha$ and conclude that the BACCs differ significantly. If the P-value is $>= \alpha$ we cannot discard the null hypothesis. To account for multiple testing, we use Bonferroni correction by multiplying each P-value by the number of tests, i.e., five tests on DNA data and nine tests on protein data.

The resulting P-values for DNA data are reported in Table S7. Here, we compared the BACC obtained from a data collection of an evolutionary model to the BACC from the data collection of the next more complex model, e.g. we compared the BACCs of JC to the BACCs of HKY.

For the GBTs, we observe a significant decrease in prediction accuracy comparing the HKY and GTR models, as well as for comparing the GTR to the GTR+G model and when comparing the two indel models (GTR+G+I+mimick and GTR+G+I+sparta).

For the CNNs, all tests were significant except for the test on the GTR+G-model and GTR+G+I-model, as well as the test comparing the indel models (GTR+G+I+mimick and GTR+G+I+sparta).

| Comparisons | JC, HKY | HKY, GTR | GTR, GTR+G | GTR+G, GTR+G+I | mimick, sparta |
|---|---|---|---|---|---|
| P-value GBT | 1.0 | 0.10 | 0.0 | 1.0 | 0.0 |
| P-value CNN | 0.03 | 3E-13 | 0.04 | 1.0 | 1.0 |

Table S7: P-values of unpaired two-samples t-tests on BACCs of the GBT and CNN on the DNA data collections with Bonferroni correction.

For protein data, we conducted tests comparing the results of each evolutionary model to the next more complex model, and all models that involve mixtures of stationary distributions (i.e., C60 and S256).

As table S8 shows, all tests comparing site homogenous models are significant for the GBTs indicating that the respective BACCs differ significantly. We observe similar results comparing the BACCs of the LG+S256 data collection with the LG+C60, LG+S256+G4, and the LG+S256+GC data collections.

| t-test P-values (GBT) | | | |
|---|---|---|---|
| | Poisson | WAG | LG |
| WAG | 0.0009 | | |
| LG | | 0.0012 | |
| LG+C60 | | | 0.0 |
| | LG+C60 | LG+S256 | LG+S256+G4 |
| LG+S256 | 0.0 | | |
| LG+S256+G4 | 1.0 | 0.0 | |
| LG+S256+GC | 1.0 | 0.0 | 1.0 |

Table S8: P-values of unpaired two-samples t-tests on BACCs of the GBTs for the protein data collections. The top half shows all tests that include a site homogeneous model. The bottom half of the table shows all combinations of evolutionary models that include C60 or S256.

For the CNNs, tests comparing site homogeneous models and LG and LG+C60 models are significant, as shown in the upper half of table S9. However, for all combinations of site heterogeneous models, the null hypothesis that their mean BACC across folds is equal cannot be rejected (see lower half of table S9).

| t-test P-values (CNN) | | | |
|---|---|---|---|
| | Poisson | WAG | LG |
| WAG | 1E-10 | | |
| LG | | 5E-07 | |
| LG+C60 | | | 2E-13 |
| | LG+C60 | LG+S256 | LG+S256+G4 |
| LG+S256 | 1.0 | | |
| LG+S256+G4 | 1.0 | 0.38 | |
| LG+S256+GC | 1.0 | 1.0 | 1.0 |

Table S9: P-values of unpaired two-samples t-tests on BACCs of the CNNs for the protein data collections with Bonferroni correction and $\alpha = 0.05$. The top half shows all tests that include a site homogeneous model. The bottom half of the table shows all combinations of evolutionary models that include C60 or S256.

### 6 TreeBASE Data

TreeBASE [8] is a heterogeneous database of published alignments. TreeBASE does not impose a limit on the kind of data that can be submitted (neither in terms of species nor in terms of types of genes/loci of the data). Determining the exact composition of underlying loci of all TreeBASE data would require parsing all associated publications of all obtained MSAs, which is not feasible in the scope of this work. Yet, to ensure that the structure detected by our GBTs in empirical data collection from TreeBASE is not due to the genetic code, we computed the number of stop codons in all genes in the data. To further ensure that we are not missing stop codons due to a shift in the reading frame, we computed the number of stop codons without shift, with one, as well as with two shifts in each sequence. Figure S17 shows the frequency of number of stop codons for each of the three computations. We do not observe an excess of data with zero or one stop codons. Note that figure Figure S17 depicts the count of stop codons only up to the 95% percentile for better visualization.

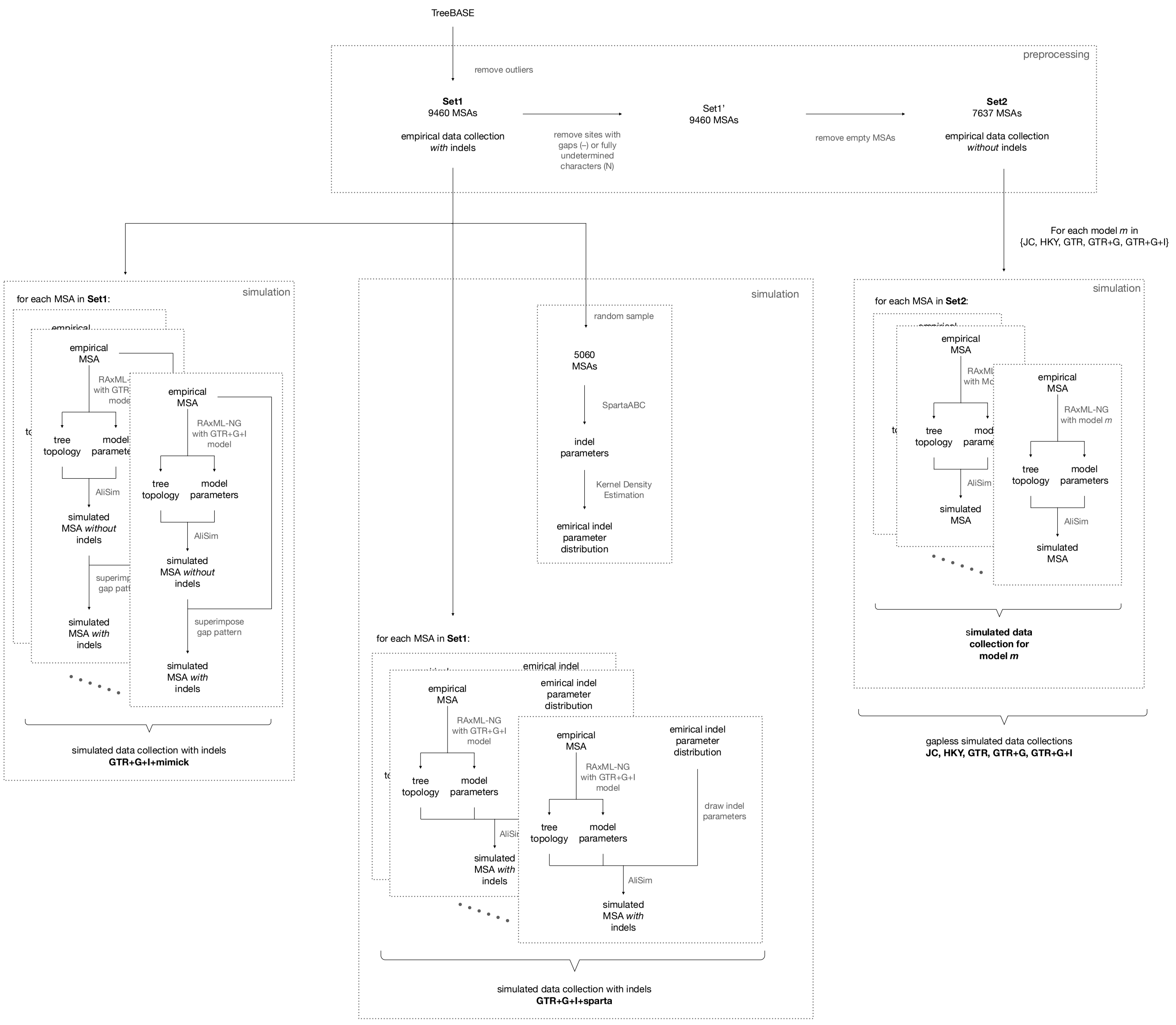

Figure S1: Schematic overview of the DNA alignment simulations.

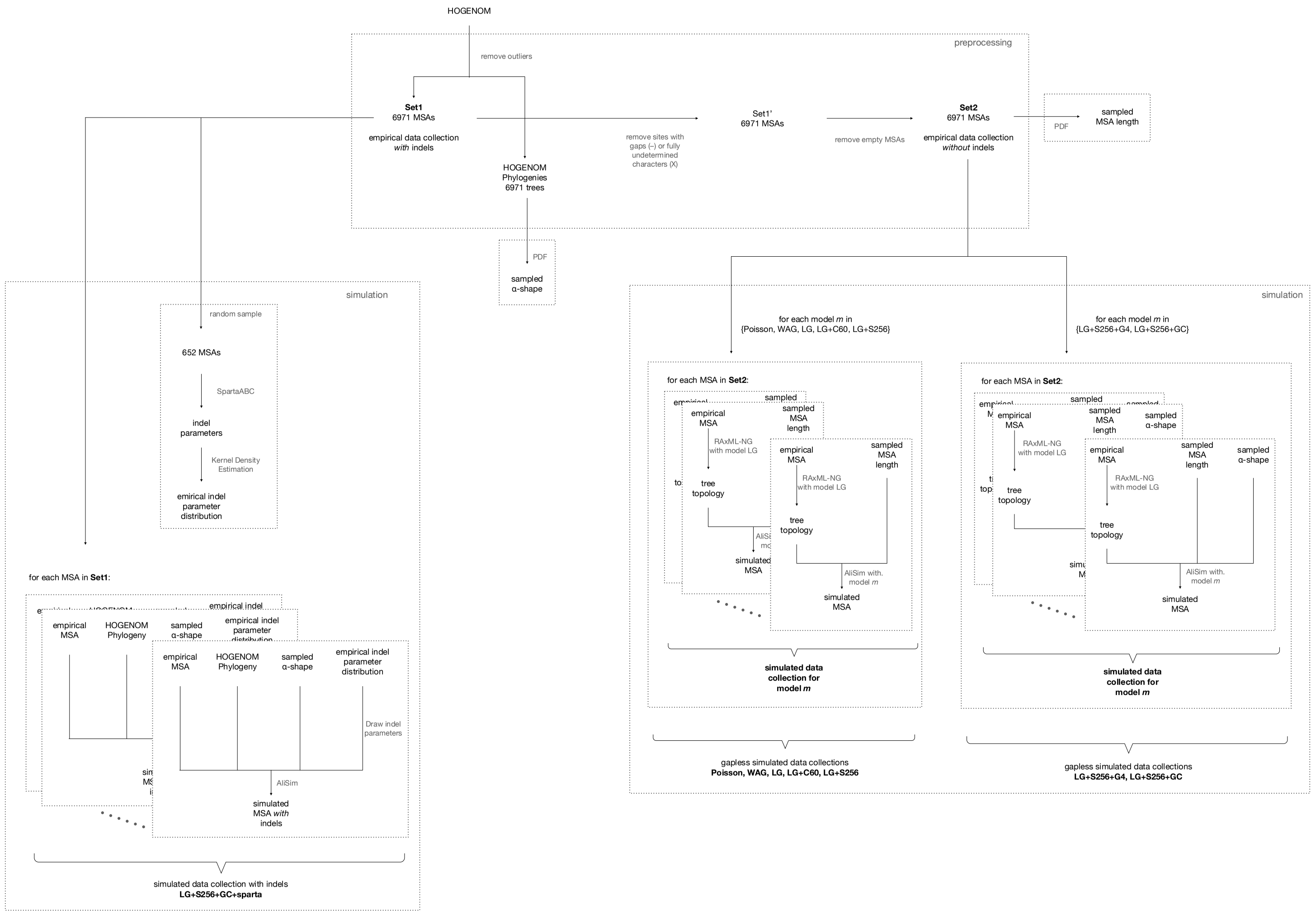

Figure S2: Schematic overview of the protein alignment simulations.

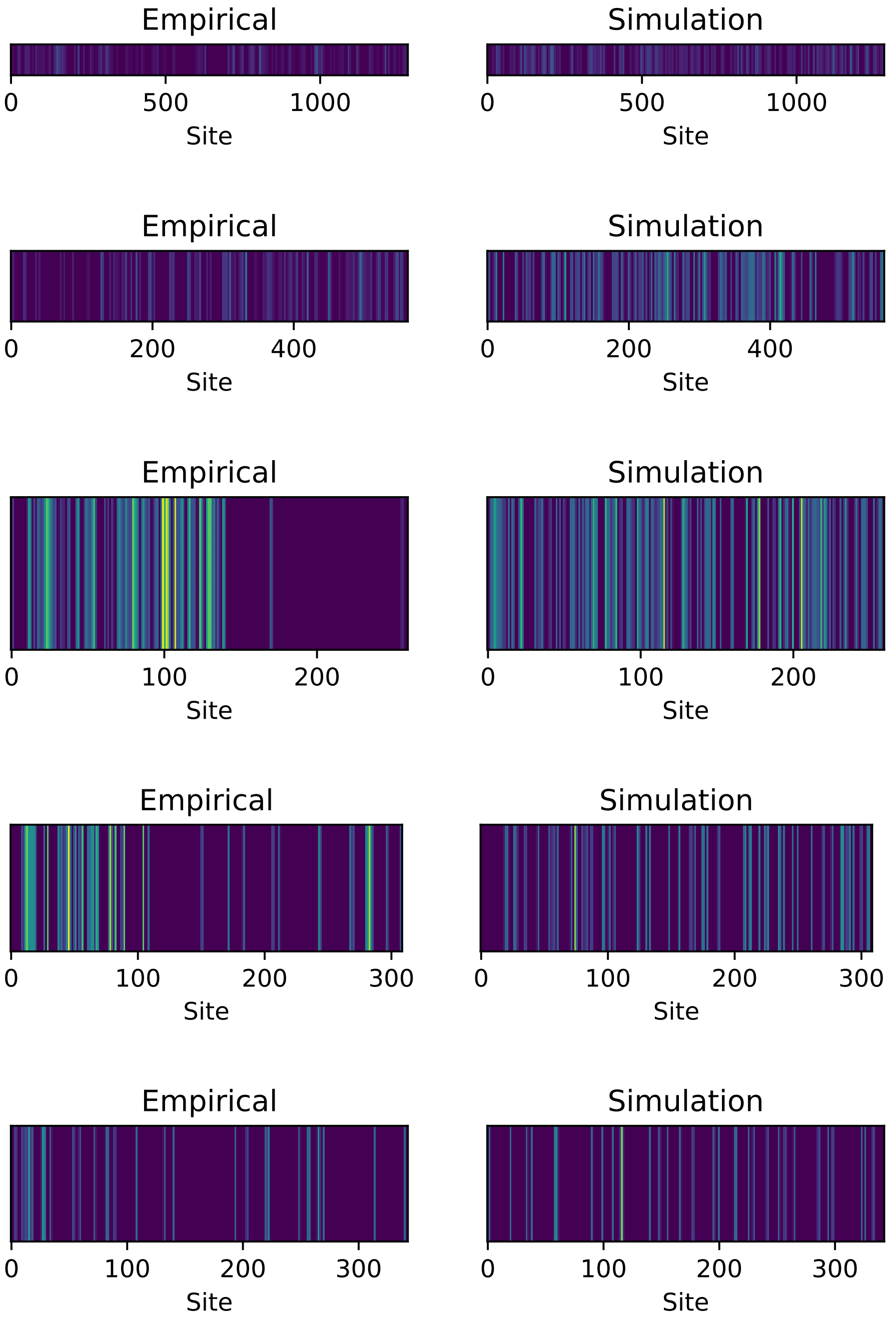

Figure S11: Visualized substitution rates for gapless empirical DNA MSA, and gapless simulated MSA generated based on the inferred tree and estimated evolutionary model parameters of the left MSA under the GTR model, for MSAs with IDs 40-44. The x-axis denotes the alignment site index. A brighter color denotes a higher number of substitutions.

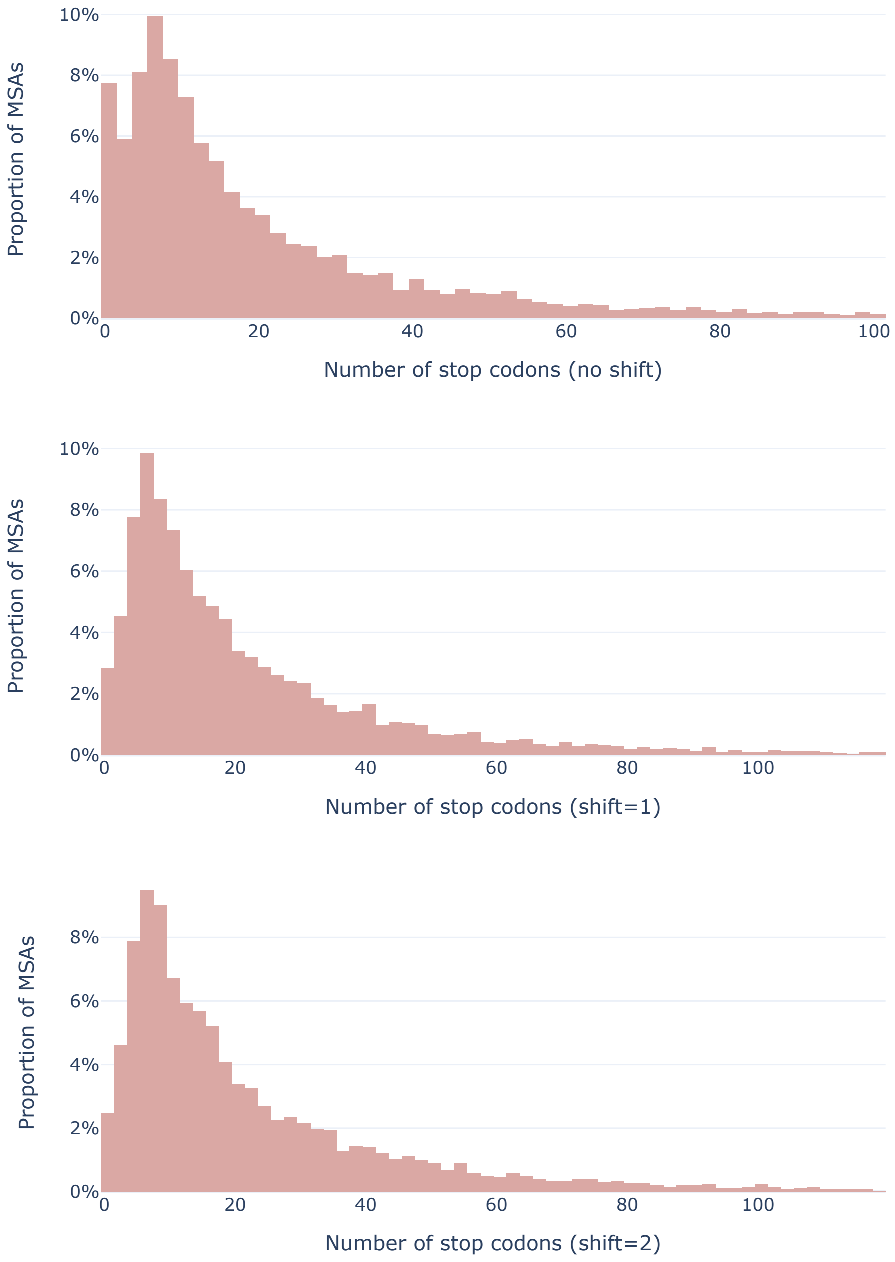

Figure S17: Number of stop codons in all translated DNA sequences of all MSAs of the empirical TreeBASE data collection.
